# A presynaptic source drives differing levels of surround suppression in two mouse retinal ganglion cell types

**DOI:** 10.1101/2022.11.12.516278

**Authors:** David Swygart, Wan-Qing Yu, Shunsuke Takeuchi, Rachel R. O.L. Wong, Gregory W. Schwartz

## Abstract

In early sensory systems, cell-type diversity generally increases from the periphery into the brain, resulting in a greater heterogeneity of responses to the same stimuli. Surround suppression is a canonical visual computation that begins within the retina and is found at varying levels across retinal ganglion cell types. Our results show that divergence in the level of surround suppression occurs subcellularly, at bipolar cell synapses. Using single-cell electrophysiology and serial block-face scanning electron microscopy, we show that two retinal ganglion cell types exhibit very different levels of surround suppression even though they receive input from the same set of bipolar cell types. This divergence of the bipolar cell signal occurs through synapse-specific regulation by amacrine cells at the scale of tens of microns. These findings indicate that each synapse of a single bipolar cell can carry a unique visual signal, expanding the number of possible functional channels at the earliest stages of visual processing.

## Introduction

Visual processing is already well underway in the retina. The analog luminance, contrast, and wavelength representation that begins in photoreceptors are transformed into >40 unique, behaviorally relevant channels of digital information that exit the retina via spikes in retinal ganglion cell (RGC) axons. Stratification of the presynaptic bipolar cell (BC) and amacrine cell (AC) interneurons and the RGC dendrites within sublaminae of the inner plexiform layer (IPL) is an established organizing principle by which retinal circuits build feature selectivity^1–4^. Nonetheless, the number of functionally distinct RGC types exceeds their stratification diversity^1,5,6^. What circuit motifs enable RGC types with nearly identical stratification patterns to have different light responses?

Previous studies have identified contributions to functional divergence from precise wiring specificity even within the same IPL sublamina^7^ or from differences in intrinsic properties of the RGCs^8,9^. Here, we examine such an example where two RGC types receive the same set of excitatory inputs but exhibit functionally distinct output signals. We isolate the circuit location at which their functions diverge, and surprisingly, it is at the level of BC output synapses, despite the commonly held view of BCs as electrically compact neurons that constitute a single information channel.

We compared two RGC types in the mouse (Pix_ON_ and ON alpha) that share very similar patterns of IPL stratification but show a striking difference in feature selectivity. The visual feature that we investigated is surround suppression: one of the oldest and best-studied visual computations^10^. The first recordings of the receptive fields (RFs) of mammalian RGCs showed a center region that was antagonized by the surrounding region, resulting in weaker signals to large stimuli than stimuli covering only the RF center^11^. Over the many decades of work that followed, it has become clear that surround suppression is not computed by a single mechanism, but instead differs by species and cell types and can arise at multiple locations in the retina^10^. We sought to identify the circuit locations at which surround suppression is computed in Pix_ON_ RGCs, where it is particularly prominent^12^ as compared to ON alpha RGCs, where it is much weaker^13^.

Surround suppression has largely been considered to be driven by wiring patterns between specific cell types. However, we show that Pix_ON_ and ON alpha RGCs have very similar circuit connectivity, particularly in their excitation, but show very different surround suppression. We find that these differences in suppression are inherited from differences in the RGC presynaptic excitatory drive, suggesting that this computation occurs at the subcellular level. These findings reveal a new location for the computation of a classical receptive field property. More generally, they suggest that subcellular computation imparts neural circuits with even more capacity for functional divergence than can be inferred from their synaptic wiring diagrams.

## Results

### The Pix_ON_ RGC has stronger surround suppression than the ON alpha RGC

We identified Pix_ON_ and ON alpha RGCs by their unique morphology and light responses^6,12,13^. These two RGC types have large dendritic arbors that primarily stratify in sublamina 5 of the IPL and exhibit ON-sustained light responses (**Fig. 1a,b**). Despite their many similarities, the Pix_ON_ and ON alpha have been shown to correspond to two unique cell types^6,12,14,15^. Morphological characteristics, such as soma size and arbor complexity, do differ between the two cell types, and ON alpha but not Pix_ON_ RGCs are SMI-32 immunoreactive (**Supplementary Fig. 1** and [ref. ^16^]). The Pix_ON_ and ON alpha RGC types both exhibit weak intrinsic light responses and correspond to the M5 and M4 intrinsically photosensitive RGC types, respectively (**Supplementary Fig. 1** and [refs. ^14,17^]). Functionally, these RGC types exhibit differing excitatory, inhibitory, and spiking receptive fields (**Fig. 1c-k**).

**Fig. 1.**
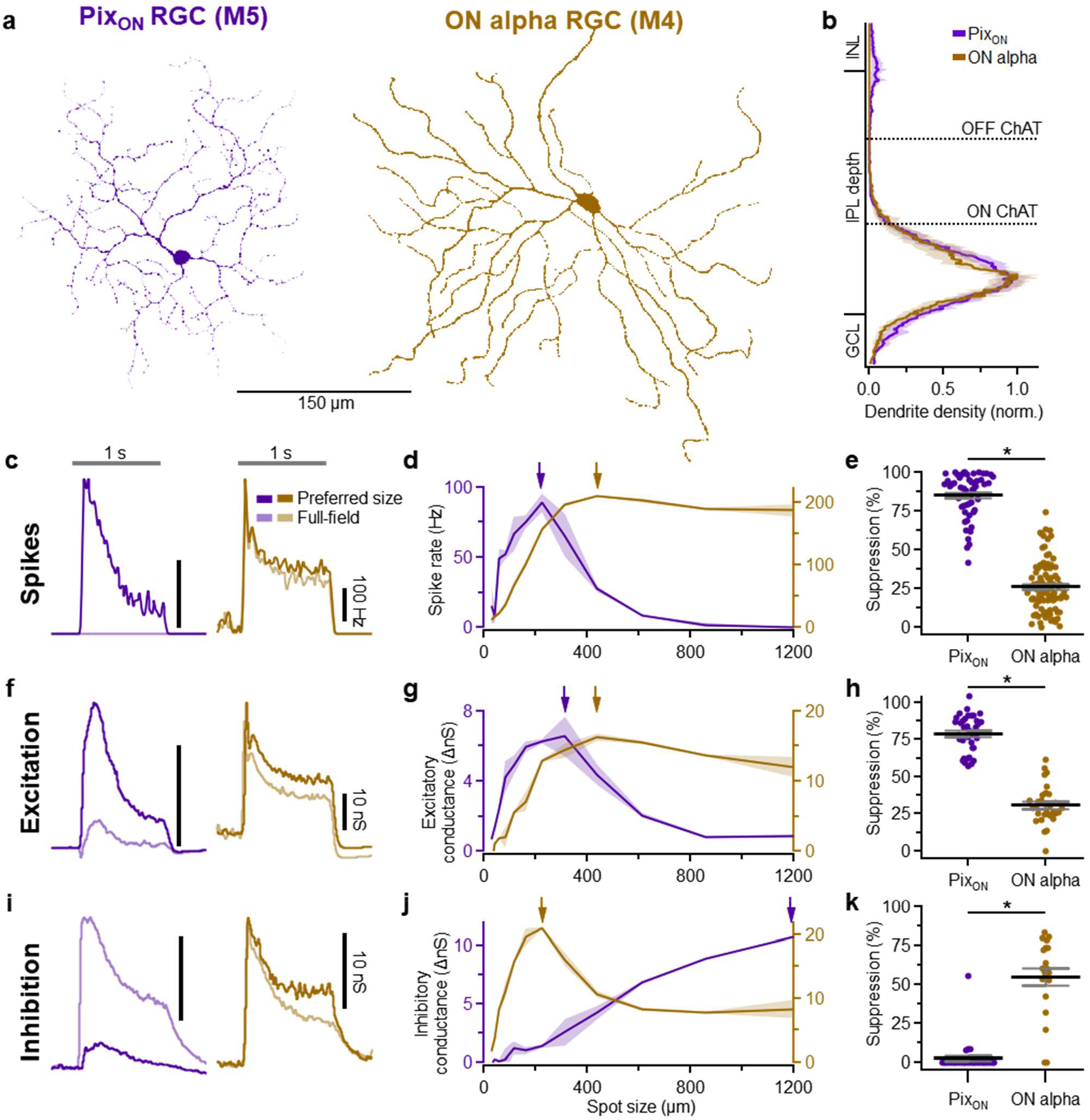
Surround suppression is stronger in Pix_ON_ RGCs than in ON alpha RGCs. **a**, *En-face* view of a Pix_ON_ (purple) and an ON alpha (brown) dendritic arbor. **b**, Dendritic stratification of Pix_ON_ (n=19) and ON alpha (n=10) RGCs within the inner nuclear layer (INL), inner plexiform layer (IPL), and ganglion cell layer (GCL). Dotted lines refer to the ON and OFF choline acetyltransferase (ChAT) bands used to determine stratification. **c**, Example peristimulus time histograms to preferred size and full-field light spot stimuli. The gray horizontal bar indicates the 1-second presentation of the 250 R*/rod/s spot stimulus from a background luminance of ∼0.3 R*/rod/s. **d**, Example spike rate across a range of spot sizes. Arrows indicate the preferred spot size. **e**, Surround suppression of spiking response for PixON (n=55) and ON alpha (n=90) RGCs. Dots indicate data from individual cells. Bar plots indicate average ± s.e.m., *P<0.05, Welch’s t-test. **f-h**, Same as **c-e** but measuring excitatory conductances via whole-cell voltage clamp configuration. **h**, PixON (n=37) and ON alpha (n=31). **i-k**, Same as **c-e** but measuring inhibitory conductances via whole-cell voltage clamp configuration. **k**, PixON (n=32) and ON alpha (n=21).

The most obvious way in which the Pix_ON_ and ON alpha RGC’s receptive fields differ is in their magnitude of surround suppression. Both RGC types exhibited ON sustained spiking responses when presented with a 200 μm diameter spot of light (**Fig. 1c**). However, when presented with large stimuli (1200 μm diameter spot), the Pix_ON_ RGC’s spike response was strongly suppressed, while the ON alpha’s spike response was only weakly suppressed (**Fig. 1d,e**; Pix_ON_ suppressed 89 ± 1.8%, n=46; ON alpha suppressed 26 ± 1.8%, n=90; P<10^−47^). This difference in surround suppression between the two RGC types was present in both scotopic and photopic conditions (**Supplementary Fig. 2**) and across retinal locations (**Supplementary Fig. 3**).

To investigate if synaptic conductances could lead to the differing levels of surround suppression in these two RGC types, we voltage-clamped both cell types and recorded excitatory and inhibitory synaptic conductances across stimulus size. Previous work demonstrated that Pix_ON_ RGCs have spatially distinct regions of their receptive fields in which they receive excitation and inhibition^12^, so we took advantage of this property to confirm that voltage-clamp effectively isolated excitation and inhibition (**Supplementary Fig. 4**). The excitatory conductances of both RGC types mirrored their spike responses; the Pix_ON_ excitatory conductances showed strong surround suppression and the ON alpha excitatory conductances showed weak surround suppression (**Fig. 1h**; Pix_ON_ suppressed 79 ± 2.0%, n=37; ON alpha suppressed 30 ± 2.1%, n=30; P<10^−24^). As previously reported^12^, the Pix_ON_ inhibitory conductances were small for small spot sizes but continually increased for larger spot sizes. In contrast, the ON alpha inhibitory conductances were large for small spot sizes and moderately suppressed for larger spot sizes (**Fig. 1k**; Pix_ON_ suppressed 2.6 ± 2.8%, n=31; ON alpha suppressed 55 ± 5.3%, n=21; P<10^−8^).

### Excitatory synaptic conductances drive surround suppression

The differing levels of surround suppression between the Pix_ON_ and ON alpha RGC types could be driven by differences in synaptic conductances (e.g., excitation and inhibition, see **Fig. 1f-k**) or by differences in cell-intrinsic factors (e.g., voltage-gated channels). To independently test the contribution of synaptic conductances and cell-intrinsic factors, we used dynamic clamp to simulate previously recorded Pix_ON_ and ON alpha excitatory and inhibitory conductances in a new set of Pix_ON_ and ON alpha RGCs (**Fig. 2a**). **Figure 2b-g** shows that strong surround suppression occurred when simulating Pix_ON_ conductances in either Pix_ON_ RGCs (suppressed 99 ± 0.4%, n=4) or ON alpha RGCs (99 ± 1%, n=3). In contrast, the simulation of ON alpha conductances induced weak surround suppression in both Pix_ON_ RGCs (18 ± 0.9%, n=4) and ON alpha RGCs (23 ± 5%, n=3). These results show that the differing levels of surround suppression in the Pixon and ON alpha spiking responses are driven by their differing conductances (99% of total variance, p<10^−12^), not by cell-intrinsic factors (0.1% of total variance, p=0.22, two-way ANOVA).

**Fig. 2.**
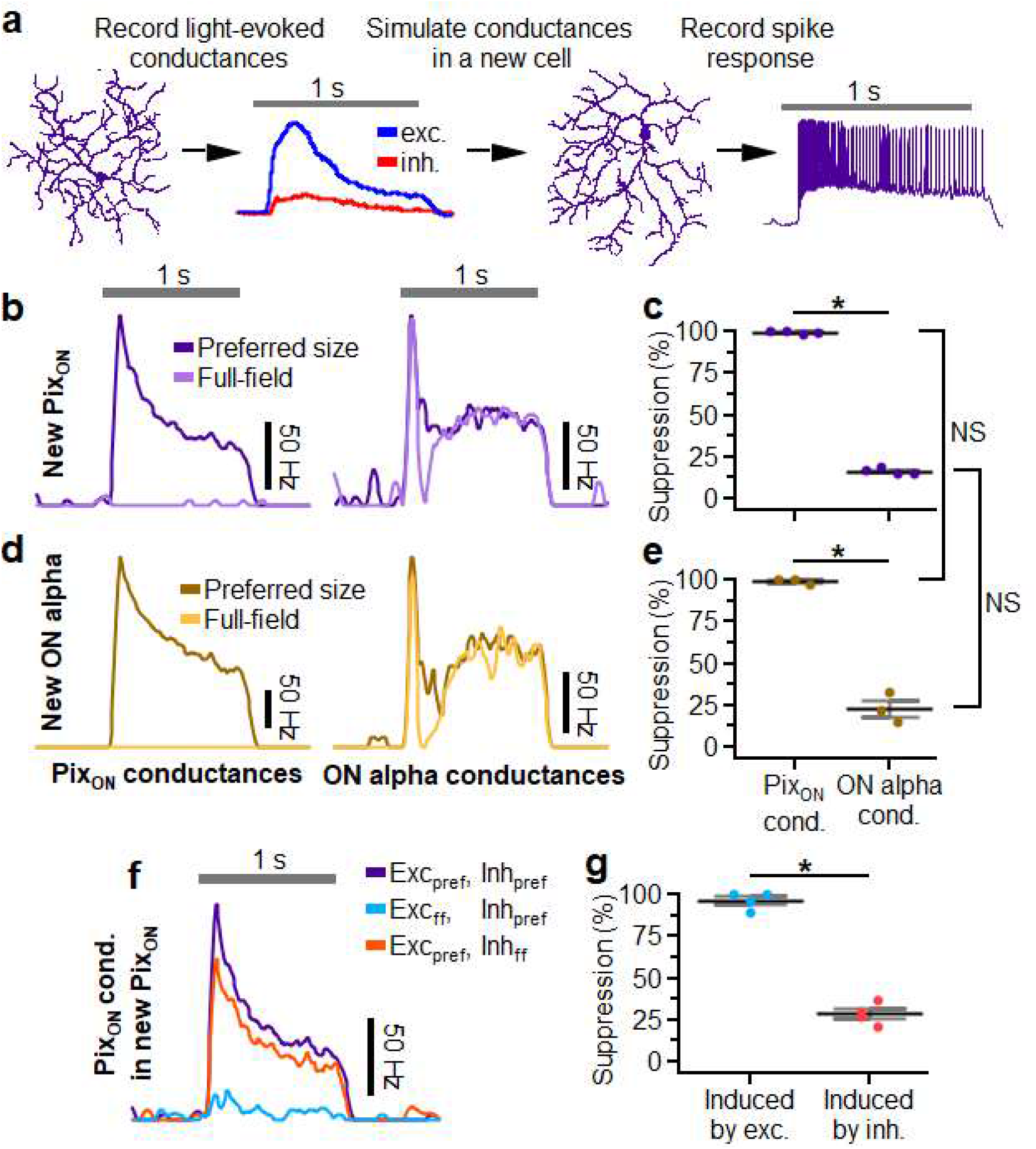
Excitatory conductances drive differing levels of surround suppression in PixON and ON alpha RGCs. **a**, Schematic illustrating dynamic clamp protocol in which previously recorded excitatory (blue) and inhibitory (red) conductances are simulated in a new RGC via current injections. **b**,**c** Example peristimulus time histogram recorded from a PixON RGC when simulating excitatory and inhibitory conductances recorded from a different PixON RGC (**b**) or an ON alpha RGC (**c**). “Preferred-size” (dark purple) indicates the maximal spiking response when simulating conductances recorded during 200, 600, and 1200 μm diameter spot stimuli. “Full-field” (light purple) indicates simulation of conductances recorded during 1200 μm spot stimuli. **d**, Surround suppression of PixON spiking responses when simulating conductances recorded from a different PixON (left) or an ON alpha (right) (n=8). **e**-**g** Same as **b**-**c** but simulating conductances within an ON alpha RGC (n=3). **h**, Example peristimulus time histogram recorded from a PixON RGC when simulating PixON conductances as to isolate the effect of full-field excitation or full-field inhibition. Purple indicates simultaneous simulation of preferred size excitation and inhibition (same as “preferred size” in **b**). Blue indicates the simulation of full-field excitation and preferred size inhibition. Orange indicates simulation of preferred size excitation and full-field inhibition. **i**, Suppression of the “preferred size” spiking response (purple trace in **h**) that is induced when switching to full-field excitation (blue trace in **h**) or full-field inhibition (orange trace in **h**) (n=4). **d**,**g**,**i** Dots indicate data from individual cells. Bar plots indicate average ± s.e.m., n=8 for all conditions, *P<0.05, Significance was determined by Two-way ANOVA for **d**,**g** and paired two-sample Student’s *t*-test for **i**.

To independently test the ability of the Pix_ON_ RGC’s excitatory versus inhibitory synaptic conductances to drive its strong surround suppression, we again utilized dynamic clamp. First, we simulated excitatory and inhibitory conductances for the preferred spot size. To test the role of excitation, we then measured how much the preferred spot spiking response was suppressed when switching to full-field excitatory conductances, but maintaining the same preferred size inhibition. Likewise, to test the role of inhibition, we measured how much the preferred spot spiking response was suppressed when simulating full-field inhibition and preferred size excitation (**Fig. 2f)**.

We found that both inhibition and excitation induced some level of surround suppression. However, full- field excitation induced significantly more surround suppression (96 ± 3%) than full-field inhibition (29 ± 3%; n=4, P=0.0002; **Fig. 2g**). These results suggest that suppression of the Pix_ON_ excitatory conductances by full-field stimuli is an important driver of surround suppression in the Pix_ON_ spiking output. Conversely, the absence of strong surround suppression of the ON alpha excitatory conductances allows the ON alpha RGC to exhibit very little surround suppression in its spiking output. Given the dominant role of excitatory conductances in dictating surround suppression of the spiking output, we next investigated sources that could cause the excitatory conductances of the Pix_ON_ and ON alpha RGCs to experience differing levels of surround suppression.

### Postsynaptic saturation or desensitization does not alter surround suppression

Differing levels of surround suppression between the Pix_ON_ and ON alpha excitatory conductances could result from differing expression of glutamate receptors in the two RGC types. If the ON alpha glutamate receptors more easily undergo saturation or desensitization, the excitatory response to the preferred size stimuli could be blunted compared to the full-field response, resulting in weaker surround suppression.

To test if glutamate receptor saturation or desensitization is necessary for the weak surround suppression in ON alpha RGCs, we measured surround suppression during bath application of subsaturating concentrations of either a weak glutamate receptor antagonist (700 nM kynurenic acid) or a strong glutamate receptor antagonist (300 nM NBQX). While both kynurenic acid (Kyn) and NBQX are expected to decrease the magnitude of the excitatory conductances, only the rapidly dissociating Kyn is expected to reduce glutamate receptor desensitization and saturation^18^. Excitatory conductances were significantly smaller in the presence of either Kyn or NBQX (Kyn 8 ± 7% of control, P<10^−2^; NBQX 20 ± 1% of control, P<10^−3^; n=3; **Fig. 3a-c**). However, surround suppression was not stronger in the presence of Kyn compared to NBQX (Kyn suppressed 12 ± 6%; NBQX suppressed 17 ± 1.9%; n=3, P=0.8; **Fig. 3c-e**). These results suggest that neither glutamate receptor saturation nor desensitization is responsible for the weak surround suppression observed in ON alpha excitatory conductances.

**Fig. 3.**
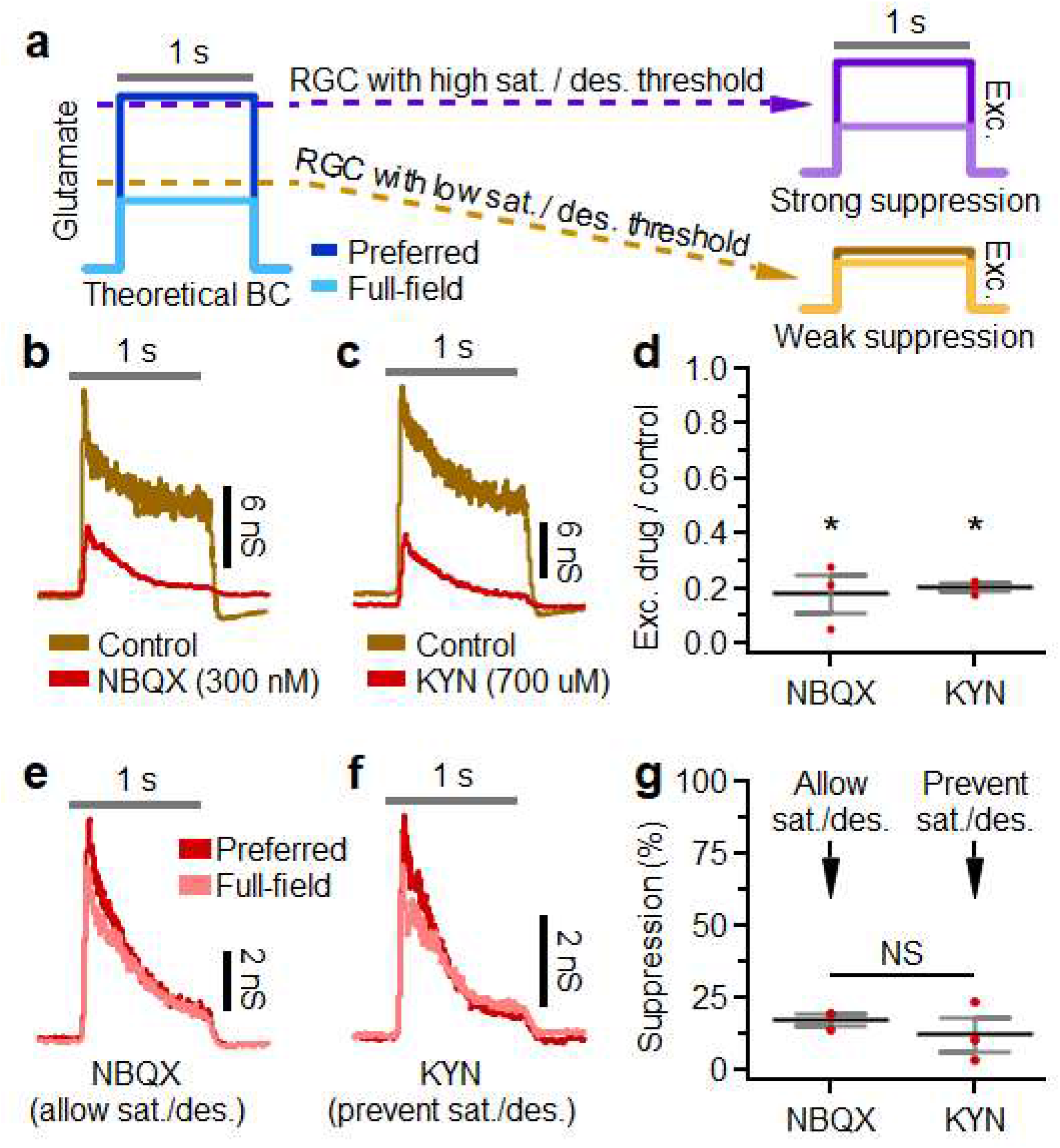
Weak surround suppression of ON alpha excitatory conductances does not depend on glutamate receptor saturation or desensitization. **a**, Example ON alpha excitatory conductances evoked by a preferred spot size in control conditions (brown) or during subsaturating bath application of NBQX (red). **b**, Same as **a**, but red indicates bath application of kynurenic acid (KYN). **c**, Proportion of ON alpha excitatory response evoked in NBQX (n=3) or KYN (n=3) compared to control conditions. **d**, ON alpha excitatory conductances evoked by a preferred (red) or full-field (pink) spot size during bath application of NBQX. **e**, Same as **d**, but during bath application of KYN. **f**, Surround suppression of ON alpha excitatory conductances in the presence of NBQX (n=3) or KYN (n=3). **a**,**b**,**d**,**e**, Gray horizontal bar indicates a 1-second presentation of the stimulus. **c**,**f**, Dots indicate data from individual cells. Bar plots indicate average ± s.e.m., *P<0.05, paired two-sample Student’s *t*-test.

Since bath application of pharmacological agents can have unintended effects that complicate interpretation, we also tested for differential saturation of the excitatory currents to Pix_ON_ and ON alpha RGCs with a different, non-pharmacological approach. We stimulated both RGC types with a range of contrast steps. The two RGC types showed similar excitatory contrast response functions, and neither cell type experienced saturation at 100% contrast (**Supplementary Fig. 5**). Together, these results suggest that the difference in excitation to Pix_ON_ and ON alpha RGCs that drives their difference in surround suppression is not a result of differential saturation or desensitization of postsynaptic glutamate receptors.

### Surround suppression in Pix_ON_ and ON alphas is accurately predicted from differing bipolar cell input but not differing RGC dendritic arbors

Having demonstrated that glutamate receptor saturation or desensitization is not the source of functionally distinct excitation in Pix_ON_ vs. ON alpha RGCs, we shifted our investigation upstream to the presynaptic BC subunits that drive excitation. An RGC’s excitatory receptive field is composed of BC subunits sampled across its dendritic arbor, with each of these BC subunits activated according to its own receptive field. The differing levels of surround suppression between the Pix_ON_ and ON alpha excitatory conductances could occur if their BC subunits had differing receptive fields, such as Pix_ON_ BC subunits exhibiting stronger surrounds. Alternatively, the differing levels of surround suppression between the Pix_ON_ and ON alpha excitatory conductances could be driven by differences in their dendritic arbors, resulting in a different spatial sampling of their BC subunits. Although Pix_ON_ and ON alpha dendritic arbors were similar in many respects, there were differences between the two RGC types, such as the slightly larger dendritic field size of the ON alpha (**Supplementary Fig. 1d**), which could cancel out BC subunit inhibitory surrounds over a larger area, resulting in decreased surround suppression of the ON alpha excitatory conductances.

To investigate how these two variables might influence surround suppression of Pix_ON_ and ON alpha excitation, we modeled an RGC’s light-evoked excitatory conductances as the summation of BC subunits sampled across its dendritic arbor (**Supplementary Fig. 6a**). We estimated the receptive field properties of these BC subunits by supplying the model with Pix_ON_ or ON alpha dendritic skeletons and then optimizing the BC’s center-to-surround ratio (CSR) and receptive field surround size (σ_s_) so that the model output best replicated the cell’s experimentally measured excitatory response across spot sizes.

Fitting the BC receptive fields to Pix_ON_ RGCs resulted in a smaller CSR (1.1 ± 0.01, median ± median absolute deviation) and a smaller σ_s_ (75 ± 5 μm) than when fitting to ON alpha RGCs (CSR = 1.8 ± 0.2, σ_s_ = 100 ± 20 μm; **Supplementary Fig. 6c,f**). Encouragingly, these receptive field properties enabled the model to accurately reproduce the experimentally measured Pix_ON_ and ON alpha excitatory responses (**Supplementary Fig. 6b,e**) and are within the range of those estimated from ON BCs glutamate signals^19,20^.

When cross-validating these BC receptive fields, the model more accurately predicted surround suppression when testing against RGCs of the same type to which the receptive field parameters were fit but failed to accurately predict surround suppression of the opposite cell type (**Supplementary Fig. 6d,g**).

When fit to Pix_ON_ RGCs, the model underestimated surround suppression in new Pix_ON_ RGCs by only 5 ± 3% (mean ± s.e.m, n=14) but overestimated surround suppression in ON alpha RGCs by 42 ± 3% (n=8). Conversely, when fit to ON alpha RGCs, the model underestimated surround suppression in new ON alpha RGCs by only 0.8 ± 4% (n=8) but underestimated surround suppression in Pix_ON_ RGCs by 49 ± 3% (n=14).

These results suggest that surround suppression of Pix_ON_ and ON alpha excitatory conductances can be well-described by input from BC subunits with reasonable RF profiles. However, fitting to either Pix_ON_ or ON alpha RGCs did not result in converging BC RF properties and poorly predicted responses in the opposite cell type, suggesting that differing BC RF properties might be required and RGC arbors cannot explain the observed differences between Pix_ON_ and ON alpha RGCs.

To directly test if any BC RF could enable the RGC dendritic arbors to accurately predict both the Pix_ON_ and ON alpha excitatory responses, we simultaneously fit a single BC RF to both RGC types. This resulted in a BC RF that provided a poorer fit to both RGC types and whose surround size was much smaller than previously reported for BC glutamate release (CSR = 1.1 ± 0.04, σ_s_ = 44 ± 7 μm, **Supplementary Fig. 6h,i**^19,20^). When cross-validating against new Pix_ON_ and ON alpha RGCs, this BC RF underestimated surround suppression for Pix_ON_ RGCs by 29 ± 4% and overestimated surround suppression for ON alpha RGCs by 14 ± 3% (**Supplementary Fig. 6j**).

Together, these results suggest that BC subunits with different receptive field properties are capable of producing the surround suppression observed in the Pix_ON_ and ON alpha excitatory conductances but differences in the dendritic arbors of Pix_ON_ and ON alphas do not appear capable of producing their differing levels of surround suppression.

### Pix_ON_ and ON alpha RGCs receive input from the same bipolar cell types

If the functionally distinct excitation in Pix_ON_ and ON alpha RGCs is driven by functionally distinct BC input, how might this difference arise? Although Pix_ON_ and ON alpha RGCs have very similar stratification profiles in the IPL where they form synapses with BCs (**Fig. 1b**), perhaps they selectively form synapses with different BC types. To determine which BCs types synapse onto the Pix_ON_ and ON alpha RGCs, we carried out serial block-face scanning electron microscopy (SBFSEM) on retinal sections that contained overlapping dendritic arbor of functionally identified Pix_ON_ and ON alpha RGCs. We identified ribbon synapses onto the dendrites of both RGCs (Pix_ON_ n=86, ON alpha n=50,) and reconstructed their presynaptic BCs (**Fig. 4a-i**). SBFSEM revealed that the Pix_ON_ and ON alpha RGCs synapsed with the same BC types in similar proportions (**Fig. 4j**). Type 6 BCs provided the majority of excitatory synapses to both the Pix_ON_ RGC (60%) and the ON alpha RGC (52%). The remaining synapses were provided by type 7 (Pix_ON_ = 31%, ON alpha = 46%), type 8 (Pix_ON_ = 1%, ON alpha = 2%), and type 9 (Pix_ON_ = 7%, ON alpha = 0%) BCs. The proportion of input from each of these types was not found to be significantly different between the Pix_ON_ and the ON alpha (T6 P=0.7, T7 P=0.3, T8 P=0.7, T9 P=0.2, two-proportions z-test with Holm-Bonferroni correction). This is also consistent with results from a different EM volume ^21^.

**Fig. 4.**
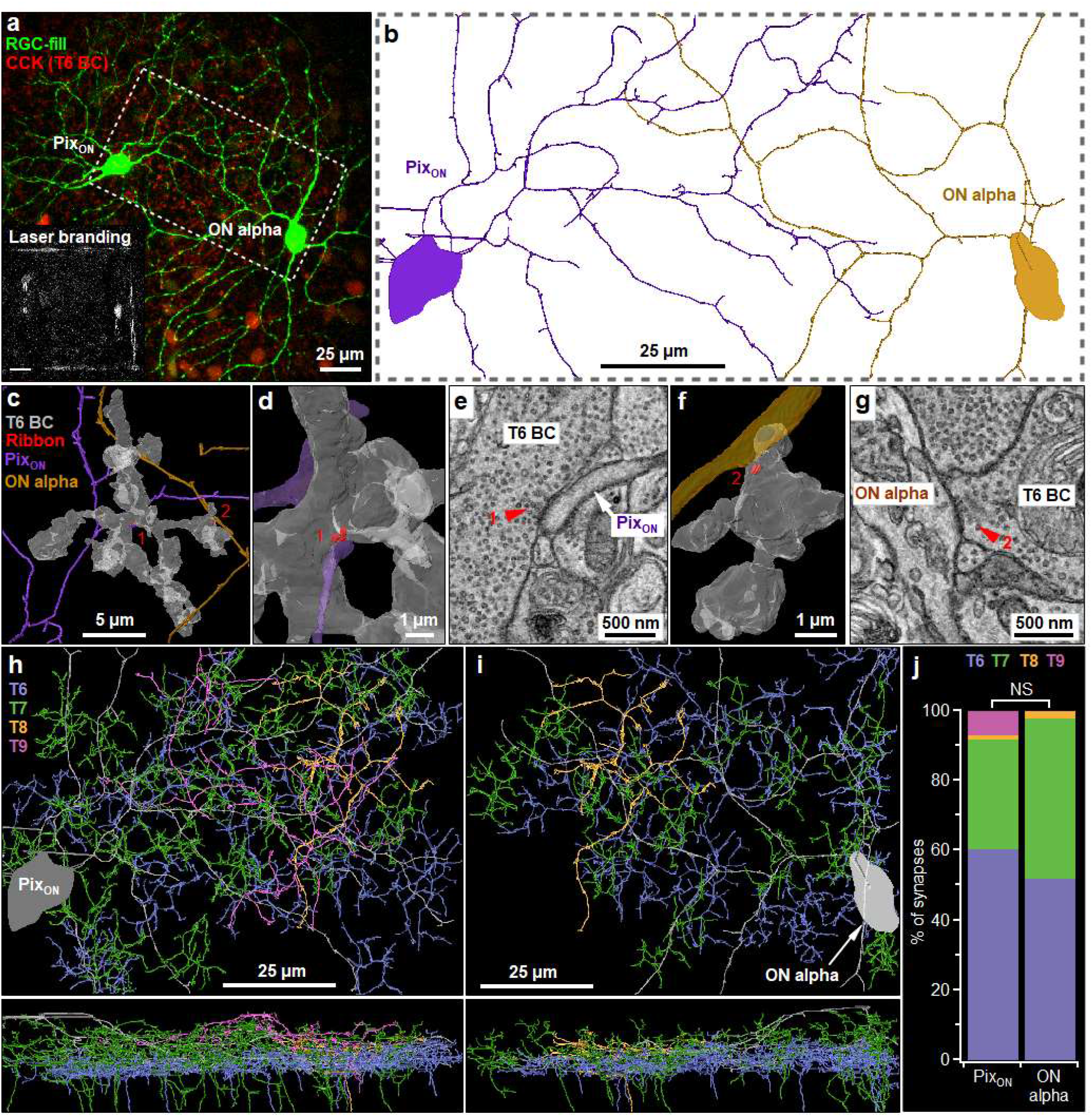
PixON and ON alpha RGCs receive excitatory input from the same bipolar cells. **a**, En-face view of filled PixON and ON alpha RGCs imaged with 2-photon microscopy in the CCK-ires-Cre/Ai14 mouse line which labels type 6 BCs (T6 BC, red). Inset shows laser burn marks used as fiducial markers during EM alignment (see methods). **b**, PixON and ON alpha SBFSEM reconstructions of the tissue volume shown in **a. c**, Example reconstruction showing a T6 BC (grey) forming ribbon synapses (red) onto a PixON dendrite (purple) and an ON alpha dendrite (brown). **d**, Reconstruction of a T6 BC ribbon synapse onto a PixON dendrite (synapse #1 from **c**). **e**, SBFSEM slice used to identify ribbon synapses (red arrow) in **d. f**,**g**, same as **d**,**e** but showing a T6 BC ribbon synapse onto an ON alpha dendrite (synapse #2 from **c**). **h**, En face (top) and cross-sectional view (bottom) of BCs types (T6-T7) presynaptic to the PixON RGC. **i**, same as **h** but for BCs presynaptic to the ON alpha RGC. **j**, The proportion of synapses formed by each bipolar cell type onto PixON (n=86) and ON alpha (n=50) RGCs. Differences in the proportion of BC type between PixON and ON alpha were not significant. P>0.05, two-proportions z-test with Holm-Bonferroni correction.

Although both RGC types received input from a similar complement of BC types, perhaps the Pix_ON_ and ON alpha RGCs form synapses with distinct subpopulations of cells within the same BC type? This hypothesis seems unlikely as we found multiple examples of individual type 6 (n=6) and type 7 (n=2) BCs that synapsed onto both the Pix_ON_ RGC and the ON alpha RGC (**Fig. 4c**). Since our SBFSEM reconstruction only covered a small area (80 × 150 μm), we sought an additional method to investigate BC input across the entire dendritic arbor of Pix_ON_ and ON alpha RGCs. To do this, we filled Pix_ON_ and ON alpha RGCs with Neurobiotin in a mouse line that fluorescently labels type 6 BCs (CCK-ires-Cre/Ai14 [refs.^22,23^]). We then used antibodies to fluorescently label an excitatory postsynaptic scaffolding protein present at excitatory synapses (PSD95^24^, **Supplementary Fig. 7a**). After confocal imaging the entire dendritic volume, we identified which PSD95-labeled synapses within the RGC dendrite were apposed to type 6 BCs (**Supplementary Fig. 7b**). In agreement with our SBFSEM results, we found that a majority of PSD95-labeled synapses were apposed to type 6 BCs for both Pix_ON_ RGCs (61 ± 2%, n=3) and ON alpha RGCs (72 ± 3%, n=2; **Supplementary Fig. 7c**) and these proportions did not significantly differ across dendritic eccentricity for either RGC type (**Supplementary Fig. 7d**, Pix_ON_ P=0.25, ON alpha P=0.17, Kolmogorov-Smirnov test).

### Amacrine cells regulate the bipolar cell terminal

If the same BCs drive excitatory conductances in both the Pix_ON_ and the ON alpha RGC types, then why is surround suppression different between the Pix_ON_ and ON alpha excitatory conductances? Perhaps surround suppression is generated at a subcellular level within the terminals of these BCs, allowing different output synapses of the same BC to convey either strong or weak surround suppression. Wide-field ACs are a promising candidate for generating surround suppression in BCs because wide-field spiking ACs have been shown to provide surround suppression of BC depolarization and glutamate release via GABA_C_ receptors clustered at cone BC output synapses^20,25–27^.

To examine the role of presynaptic inhibition by spiking ACs in generating surround suppression in Pix_ON_ RGCs, we measured Pix_ON_ excitatory conductances in the presence of GABA and glycine receptor antagonists (**Fig 5a,b**). Surround suppression was significantly decreased in the presence of a GABA_C_ receptor antagonist (control 71 ± 4%; TPMPA 34 ± 4%; n=6, P<10^−3^) but was not significantly altered by the application of a GABA_A_ antagonist (control 76 ± 9%; gabazine 80 ± 10%; n=3, P=0.8), a GABA_B_ antagonist (control 77 ± 5%; saclofen 74 ± 5%; n=4, P=0.06), or a glycine receptor antagonist (control 79 ± 9%; strychnine 78 ± 9%; n=3, P=0.7). Additionally, we found that surround suppression was significantly decreased in the presence of a voltage-gated sodium channel blocker (control 85 ± 5%; TTX 43 ± 2%; n=4, P=0.003), which is expected to block spike propagation along the neurites of spiking wide-field ACs. While TPMPA and TTX each significantly reduced surround suppression of the Pix_ON_ excitatory conductances, some surround suppression remained. However, the simultaneous application of TPMPA and TTX completely abolished surround suppression (control 71 ± 4%, TPMPA + TTX 0.3 ± 0.3%, n=2) and had a greater effect than TPMPA alone (P=0.0002), or TTX alone (P=0.0002). Experiments in ON alpha RGCs showed qualitatively similar effects but decreases in surround suppression but were more difficult to measure since surround suppression of the ON alpha excitatory conductances was already weak in control conditions (**Supplementary Fig. 8**). These results suggest that the strong surround suppression observed in the Pix_ON_ excitatory conductances is driven by spiking wide-field ACs via GABA_C_ receptors at the BC terminal. While these same cells may drive what little surround suppression is present in the ON alpha excitatory conductances, they appear unable to induce the same level of surround suppression as seen in Pix_ON_ RGCs.

**Fig. 5.**
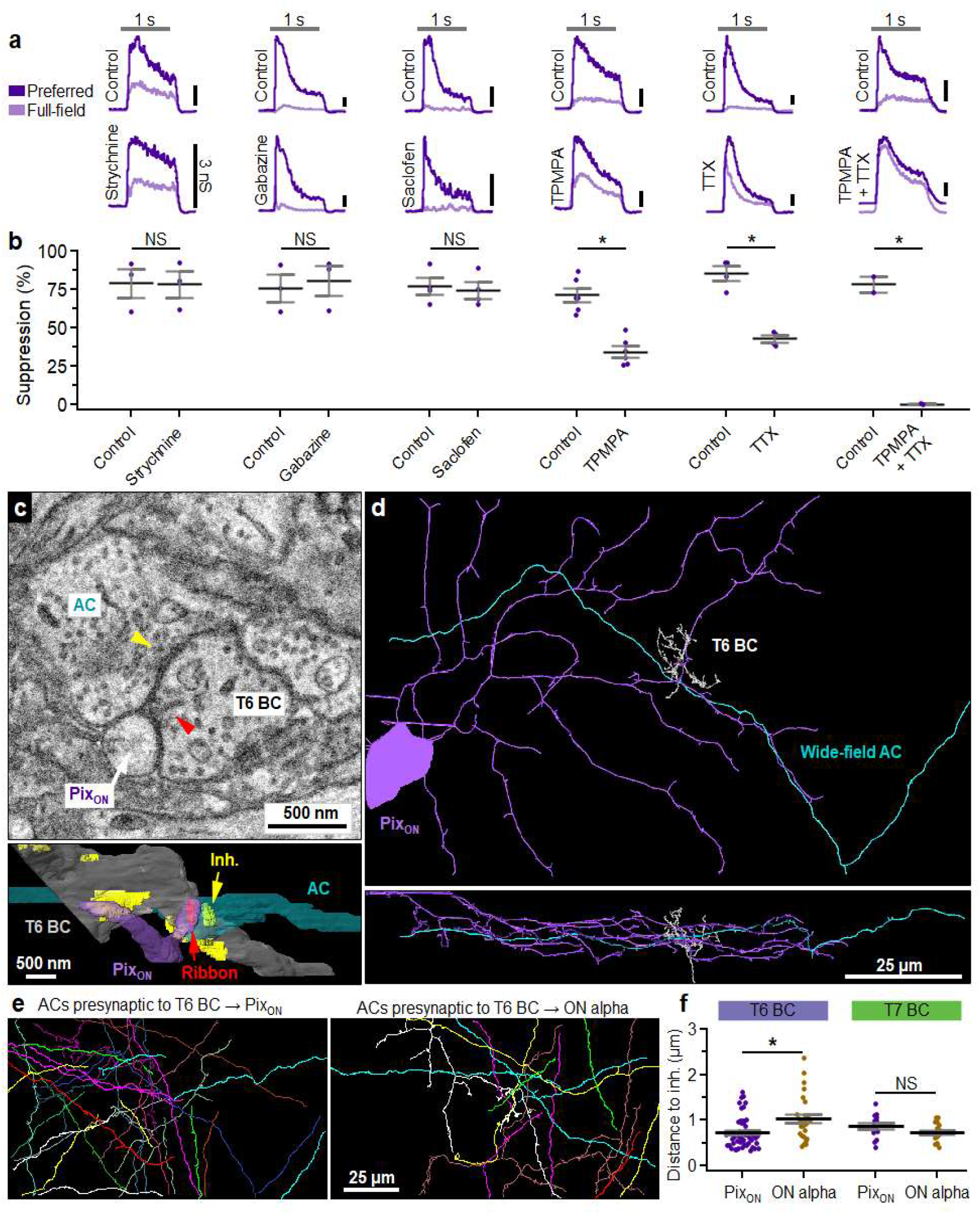
Wide-field amacrine cell regulation at the bipolar cell terminal contributes to stronger surround suppression in PixON RGCs. **a**, PixON excitatory conductances evoked before (top) and after (bottom) bath application of antagonists of glycine receptors (strychnine), GABAA receptors (gabazine), GABAB receptors (saclofen), GABAC receptors (TPMPA), or NaV channels (TTX). The gray horizontal bar indicates a 1-second presentation of the stimulus. Note: conductance trace in TPMPA+TTX was shifted down 2 nS to improve visibility. **b**, Surround suppression of excitatory conductances in control and antagonists conditions. Dots indicate data from individual cells strychnine (n=3), gabazine (n=3), saclofen (n=4), TPMPA (n=6), TTX (n=4), TPMPA+TTX (n=2). Bar plots indicate average ± s.e.m., *P<0.05, paired two-sample Student’s *t*-test. **c**, SBFSEM slice (top) and reconstruction (bottom) showing an AC neurite (cyan) forming an inhibitory synapse (yellow) onto a BC (gray) which then forms a ribbon synapse (red arrow) onto a PixON RGC dendrite (purple). **d**, A zoomed-out En face (top) and cross-sectional (bottom) view of the AC from **c. e**, Reconstruction of nearest presynaptic ACs to T6 BC-to-PixON (left) and T6-to-ON alpha (right) ribbon synapses. **f**, Distance to nearest inhibitory from T6 BC output synapses (PixON n=51, ON alpha n=26) and T7 BC output synapses (PixON n=14, ON alpha n=17). Dots indicate data from each BC-to-RGC synapse. Bar plots indicate average ± s.e.m., *P<0.05, Welch’s t-test.

While our pharmacology results suggested a role for presynaptic inhibition by spiking GABAergic ACs in generating surround suppression in Pix_ON_ RGCs, they did not offer direct evidence of differential inhibition at synapses to Pix_ON_ vs. ON alpha RGCs. To investigate presynaptic AC inhibition at a subcellular level, we reconstructed the ACs which formed output synapses onto the presynaptic BCs identified in our SBFSEM volume (from **Fig. 4**). For each type 6 and type 7 BC ribbon synapse onto the Pix_ON_ and ON alpha RGC, we identified the nearest presynaptic inhibitory site (**Fig. 5c**). Although a presynaptic inhibitory site was always found within a few microns of each BC ribbon synapse, this distance tended to be shorter for type 6 BC synapses onto the Pix_ON_ RGC (0.74 ± 0.05 μm, n=51) compared to the ON alpha RGC (1.04 ± 0.1 μm, n=26; P=0.009; **Fig. 5f**). However, this difference in presynaptic inhibitory distance was not significant for type 7 BCs (Pix_ON_ = 0.86 ± 0.07 μm, n=14; ON alpha = 0.73 ± 0.05 μm, n=17; P=0.14).

We traced the presynaptic ACs at each type 6 BC ribbon to determine if they were a likely candidate to carry inhibition from the surround. Due to the limited size of the EM reconstruction, only 60% of these ACs could be classified by field size, but all of these were identified as medium to large-field ACs (spanning >40 μm), with none of their somas contained within the reconstructed volume (**Fig. 5e**). Additionally, of the nine ACs for which we observed multiple inhibitory feedback synapses onto the type 6 BCs within the field of view, only one provided presynaptic inhibition to both a BC-Pix_ON_ and a BC-ON alpha synapses, suggesting the possibility of synapse preference based on the postsynaptic ganglion cell identity. While we could not determine the specific cell type of these wide-field ACs due to the limited size of the EM reconstruction, these results show that wide-field AC inhibition is present near each BC output synapse, but is more tightly localized at type 6 BC-Pix_ON_ synapses. This suggests that synapse-specific regulation could occur within the same BC dependent upon the identity of the postsynaptic RGC type.

### Electrotonic properties of bipolar cell terminals

Since inhibitory synapses were closer to the type 6 BC-Pix_ON_ synapses than type 6 BC-ON alpha synapses (**Fig. 5f**) and Pix_ON_ surround suppression is dependent on ionotropic GABA_C_ receptors (**Fig. 5b**), one might hypothesize that subcellular surround suppression is achieved by subcellular hyperpolarization localized to BC-Pix_ON_ output synapses. But BCs are small and so is the distance between their output synapses, bringing into doubt whether voltage could differ enough between output synapses to actually cause differing glutamate release. To investigate whether electrical compartmentalization can support functionally divergent signals from a BC, we generated a morphologically detailed NEURON cable model^28^ from an EM reconstruction of a type 6 BC, including the location of all 91 ribbon output synapses and 120 presynaptic inhibitory synapses (**Fig 6a**). Although BCs are often modeled using only passive membrane properties^29–31^, multiple studies have measured voltage-gated ion conductances from BCs which could lead to greater electrical compartmentalization^32–35^. Thus we performed all experiments in both a passive model and an active model containing L-type Ca^2+^ channels^36,37^, K_V+_ channels^38^, and HCN_2_ channels^39,40^ (see **Methods** for model details, **Table 3** for parameter values, and **Supplementary Fig. 9** for robustness tests).

**Fig. 6.**
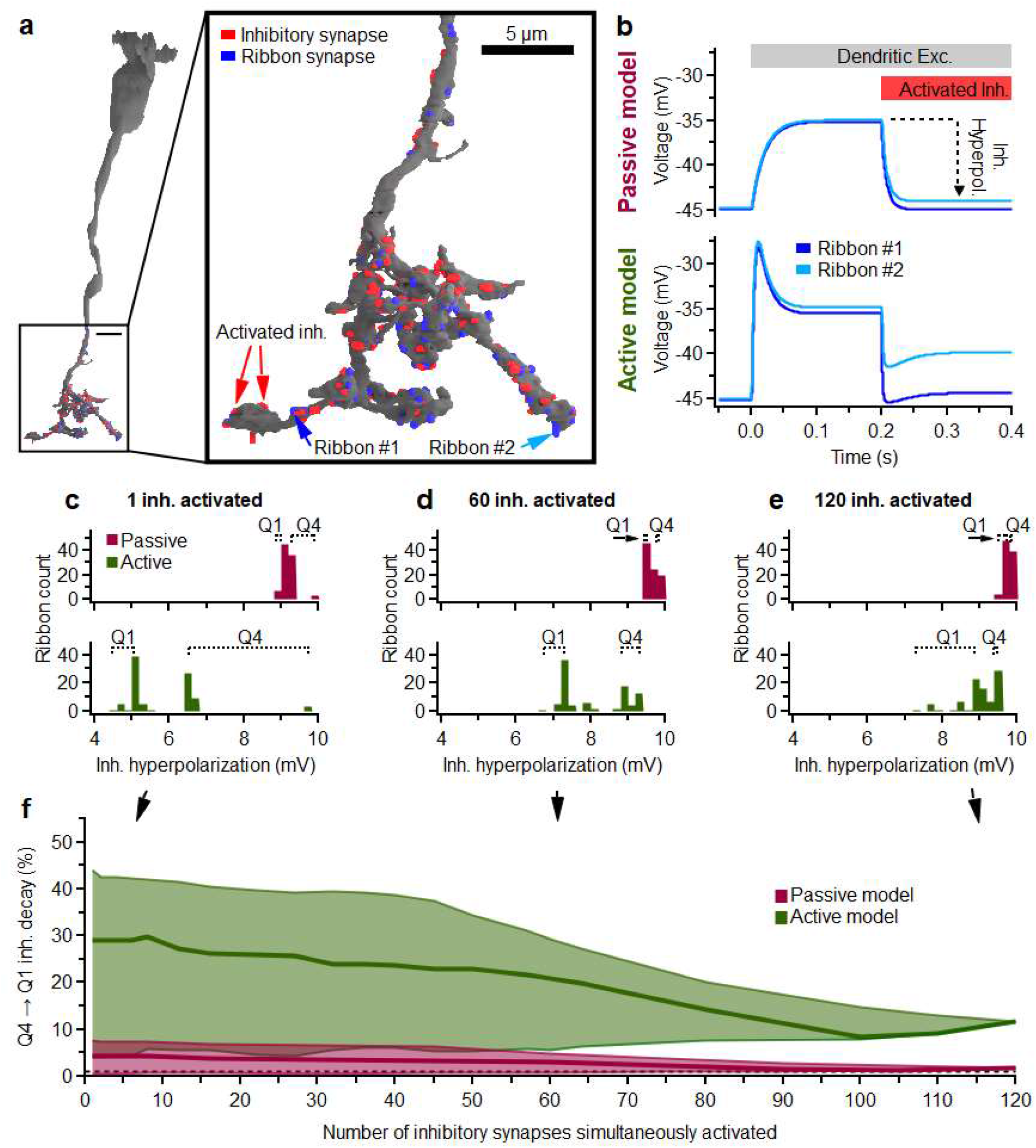
Cable model of the bipolar cell terminal. **a**, An SBFSEM reconstruction of a type 6 BC (T6 BC), including 91 ribbon output synapses (blue) and 120 inhibitory input synapses (red). **b**, Example voltage trace of ribbons in **a** (blue arrows) during a simulation experiment in which the BC is depolarized via excitatory currents at its dendrites (dendritic exc.) and then inhibitory synapses are activated (Activated inh., red arrows from **a**). *Top*, ribbon voltage measured in a passive model of the BC. *Bottom*, voltage measured in an active model of a T6 BC which includes voltage-gated channels (L-type Ca^2+^, KV+, and HCN2). **c**, Example histogram of inhibitory hyperpolarization of the ribbon synapses in the passive (*top*) and active (*bottom*) BC model when activating a single inhibitory synapse. Q1 and Q4 indicate the range of the top and bottom quartiles of ribbon hyperpolarization. **d**,**e**, Same as **c** but when simultaneously activating 60 (**d**) or 120 (**e**) inhibitory synapses. **f**, Decrement of the average inhibitory hyperpolarization of Q2 ribbons to Q1 ribbons based on the number of inhibitory synapses simultaneously activated. Thick lines indicate the median decay across all sets of inhibitory synapses activated. Shading indicates the range of inhibitory decay across all sets of activated inhibitory synapses (n=120).

**Table 1.**
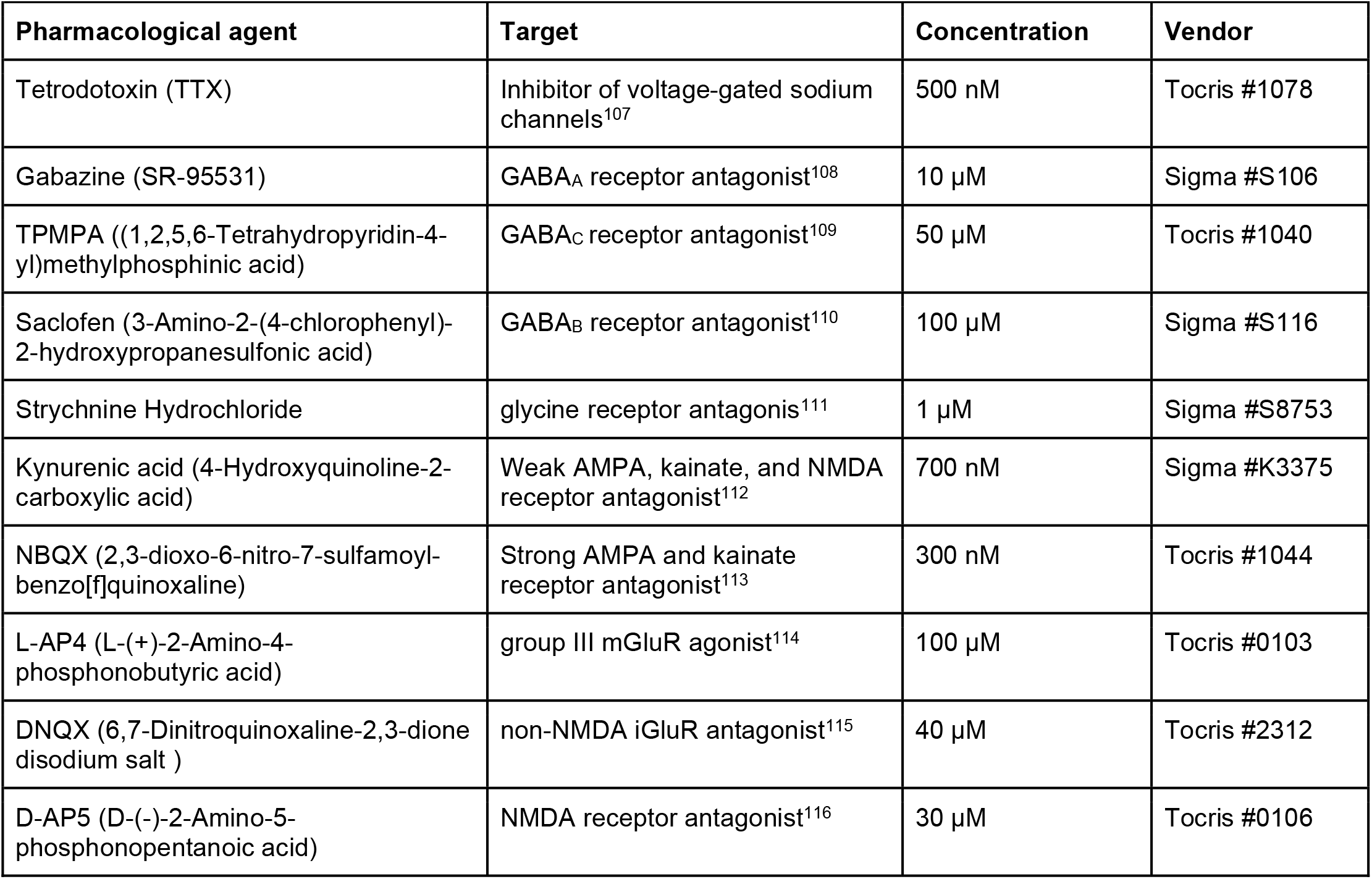
Pharmacological agents.

**Table 2.**
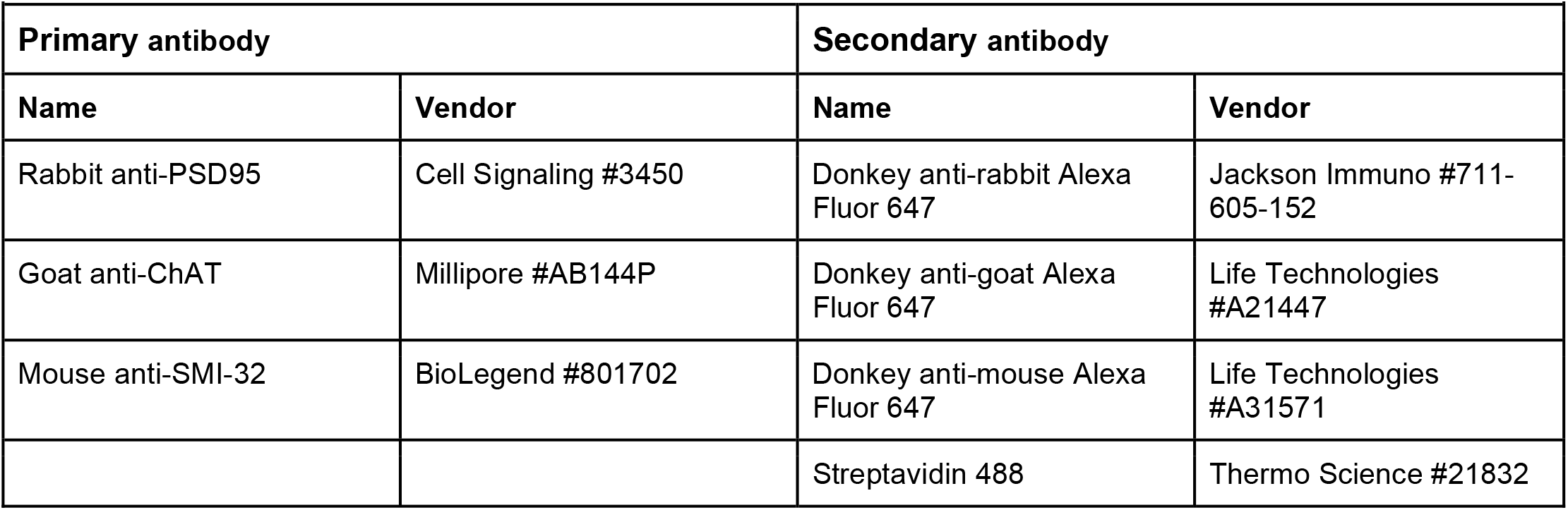
Antibodies used in immunohistochemical labeling.

**Table 3.** Key parameters of type 6 BC NEURON model.

| Table 3 \| Key parameters of type 6 BC NEURON model |  |  |
| --- | --- | --- |
| Experimental parameters | Time step | 25 μs |
|  | Temperature | 32° |
| Passive properties | Membrane capacitance | 11.8 fF/μm<sup>2</sup> [ref. <sup>29</sup>] |
| | Cytoplasmic resistivity | 13.2 mOhm-μm [ref. <sup>29</sup>] |
| | Leak conductance | 3.9 pS/μm<sup>2</sup> [ref. <sup>29</sup>] |
|  | Leak reversal potential | -60 mV [ref. <sup>29</sup>] |
| Excitation at dendrites<br>(8 synapses) | Reversal potential | 10.1 mV [ref. <sup>117</sup>] |
|  | Total conductance (passive model) | 334 pS [ref. <sup>41,118</sup>] |
|  | Total conductance (active model) | 2,600 pS [ref. <sup>41,118</sup>] |
| Inhibition at axon terminals | Reversal potential | -50.4 mV [ref. <sup>119</sup>] |
|  | Conductance (passive model) | 1,530 pS [ref. <sup>120</sup>] |
|  | Conductance (active model) | 8,000 pS [ref. <sup>120</sup>] |
| HCN2 channel<br>(restricted to axonal arbor)<br>[ref. <sup>121</sup>] | reversal potential | -23.4 mV <sup>122</sup> |
| | max conductance | 0.78 pS/μm<sup>2</sup> [ref. <sup>39,40</sup>] |
|  | Half-max of activation | -99 mV [ref. <sup>123</sup>] |
|  | Slope of activation | -6.2 mV [ref. <sup>123</sup>] |
| K<sub>v</sub><sup>+</sup> channel [ref. <sup>124</sup>] | K<sup>+</sup> reversal potential | -84 mV |
| | Max conductance | 12 pS/μm<sup>2</sup> [ref. <sup>38</sup>] |
|  | Half-max of activation | -9 mV [ref. <sup>125</sup>] |
|  | Slope of activation | 14 mV [ref. <sup>125</sup>] |
|  | Half-max of inactivation | 8 mV [ref. <sup>125</sup>] |
|  | Slope of inactivation | -9 mV [ref. <sup>125</sup>] |
| L-type Ca<sup>2+</sup> channels<br>(restricted to axonal arbor)<br>[ref. <sup>126</sup>] | Ca<sup>2+</sup> reversal potential | 18 mV [ref. <sup>37</sup>] |
| | L-type Ca<sup>2+</sup> max conductance | 1.6 pS/μm<sup>2</sup> [ref. <sup>37</sup>] |
|  | L-type Ca<sup>2+</sup> half-max activation | -32 mV [ref. <sup>37</sup>] |
|  | L-type Ca<sup>2+</sup> slope of activation | 10 mV [ref. <sup>37</sup>] |
|  | L-type Ca<sup>2+</sup> half-max inactivation | 10 mV [ref. <sup>37</sup>] |
|  | L-type Ca<sup>2+</sup> slope of inactivation | -12 mV [ref. <sup>37</sup>] |

To estimate the ability of the inhibitory sites to differentially suppress ribbon output synapses, we depolarized the BC by activating excitatory synapses on the dendrites. We then activated sets of presynaptic inhibitory synapses and measured the resulting hyperpolarization at each ribbon output synapse (**Fig 6b**). When activating a single inhibitory synapse, the magnitude of the hyperpolarization varied across the 91 ribbon output synapses (**Fig 6c**). To quantify this variation, we calculated the percent decrease of hyperpolarization from the top to the bottom quartile of ribbons. This measure of inhibitory voltage decay varied based on which inhibitory synapse was activated, but tended to be much larger for the active model compared to the passive model, with a median hyperpolarization decay of 29% in the active model and only 4% in the passive model. These data are plotted as a function of distance in **Supplementary Fig. 10** to estimate the effective electrotonic length constant of each inhibitory synapse for each model.

As it seems unlikely that the BC’s inhibitory surround is actually conveyed by a single inhibitory synapse, we also performed this same experiment while simultaneously activating multiple inhibitory synapses. To assess how the simultaneous activation of multiple inhibitory synapses impacts the spread of hyperpolarization, we repeated the sequential activation of each of the 120 inhibitory synapses but also included the simultaneous activation of the n-nearest neighbors to that synapse. As more inhibitory synapses were simultaneously activated, the range of hyperpolarizations across the ribbons decreased (**Fig 6c-e**) with a corresponding decrease in the inhibitory voltage decay between the top and bottom quartiles of ribbons (inhibitory decay of 1.8% for the passive model and 12% for the active model when activating all 120 inhibitory synapses, **Fig 6f**).

We wanted to test if the compartmentalization of inhibition seen in either the passive or active model of the BC could enable the differing levels of surround suppression measured in the Pix_ON_ and ON alpha excitatory conductances. However, this requires moving beyond a model of a single BC, as an RGC receives input from many BCs across its dendritic arbor. Thus, we combined the results of the BC cable model with the previously described BC subunit model that predicts an RGC’s excitatory response as the summation of BC receptive field subunits sampled across its dendritic arbor (**Supplementary Fig. 6a**).

If voltage decay of the BC’s inhibitory surround causes the weakened surround suppression of the ON alpha’s excitatory input compared to the Pix_ON_, then the two RGCs would have to selectively sample from the pool of ribbon output synapses based on their level of inhibitory decay. Thus, we assigned the quartile of ribbons with the greatest hyperpolarization to the Pix_ON_ and the quartile of ribbons with the least hyperpolarization to the ON alpha (**Fig. 7a**). This was an arbitrary assignment, and further research is needed to determine if this level of functionally specific synapse formation actually occurs.

**Fig. 7.**
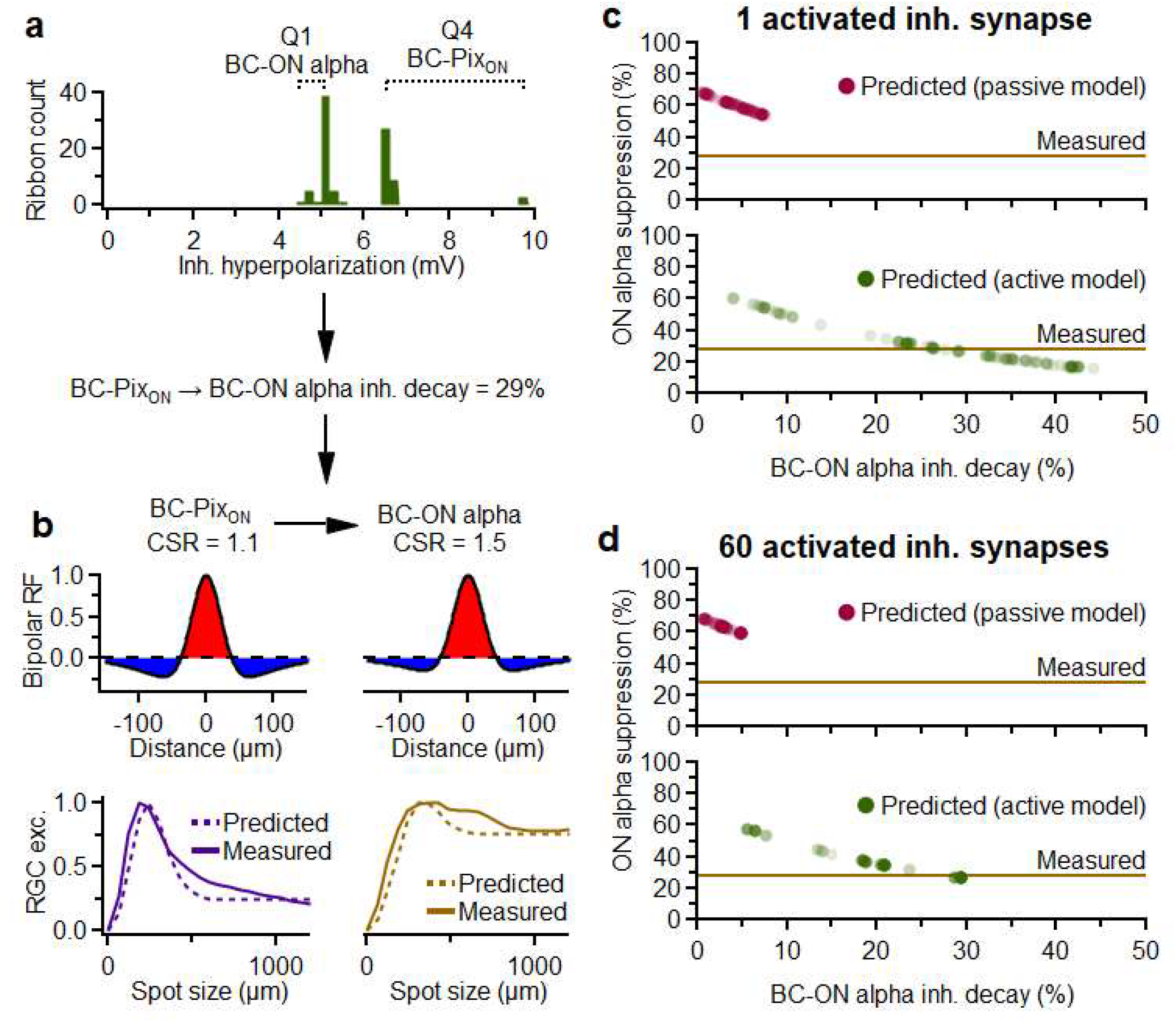
Modeling suppression of RGC excitation through electrical isolation of inhibition on the bipolar axonal arbor. **a**, Example histogram of hyperpolarization experienced by the BC ribbon output synapses upon activation of an inhibitory synapse (see **Fig. 6**). If the top (Q4) and bottom (Q1) quartiles of ribbons are assumed to synapse on PixON and ON alpha respectively, then inhibition of ON alpha BC input would be expected to decay by 29% compared to the PixON for this example. **b**, *Top*, BC center to surround ratio (CSR) and receptive fields predicted for PixON and ON alpha BC subunits given a 29% inhibitory decay (assumptions described in text). *Bottom*, Resulting model predictions (dotted lines) for surround suppression of PixON (n=14) and ON alpha (n=8) excitatory conductances according to the BC subunit model detailed in **Supplementary Fig. 6**. Solid lines indicate average experimentally measured excitatory responses for the same cells. **c**, Inhibitory voltage decay predicted for each inhibitory synapse in the passive (*Top*) or active (*Bottom*) models of the T6 BC and the resulting predictions of ON alpha surround suppression. The brown line indicated experimentally measured ON alpha surround suppression (n=8). **d**, Same as **c** but simultaneously activating sets of 60 inhibitory synapses.

If we additionally assume that voltage is linearly related to glutamate release, we can infer the relative center-to-surround ratio (CSR) of the Pix_ON_ and ON alpha’s BC subunits from the voltage decay of inhibition at their assigned ribbons (i.e. if the BC-ON alpha ribbons experience 50% inhibitory decay compared to BC-Pix_ON_ ribbons, then the ON alpha BC subunits will have CSR values twice as large as Pix_ON_ BC subunits). We fixed the Pix_ON_ BC subunit CSR value to 1.1, as it accurately predicted surround suppression for the Pix_ON_ (**Supplementary Fig. 6a)**. We then determined the ON alpha BC subunit CSR from the inhibitory voltage decay at its ribbons and predicted surround suppression of the ON alpha excitatory input using the BC subunit model (**Fig. 7a,b**).

**Figure 7c** shows that in the passive BC model, none of the individual inhibitory synapses provided inhibition with enough decay to support the decreased surround suppression measured from the ON alpha excitatory conductances. However, in the active BC model, 62 of the 120 inhibitory synapses were predicted to cause a level of surround suppression equal to or less than that measured in the ON alpha. When repeating this experiment with the simultaneous activation of half of the BC’s 120 inhibitory synapses, 41 of the 120 sets of 60 inhibitory synapses continued to predict a level of surround suppression equal to or less than the measured ON alpha surround suppression (**Fig. 7d**).

These modeling results suggest that the BC axonal arbor is not completely isopotential and that voltage gradients could feasibly contribute to localized glutamate release, particularly since a nonlinear relationship between voltage and glutamate release^41,42^ could accentuate differences between ribbons. However, a voltage gradient within the BC arbor is not absolutely required for functional divergence. We speculate in the **Discussion** about chemical sources of subcellular functional divergence at small spatial scales within the BC terminal that could work in concert with the voltage gradient or independently.

## Discussion

Our study identified the site of functional divergence between an RGC type with strong surround suppression (Pix_ON_) and a type with weak suppression (ON alpha). Contrary to the prevailing view of the functional organization of the retina and the central nervous system more generally, this site of divergence was not different types of neurons but instead nearby (<20 μm) synapses within the axon terminal of the same neuron. The capacity for functional compartmentalization at such a small spatial scale requires a new framework for information processing in the excitatory pathways of the retina. These results, along with an increasing set of similar observations throughout the brain, suggest that more detailed biophysical work on the pre-synapse is required to appreciate the computational complexity of neuronal output.

### Subcellular output divergence in the retina and the brain

In spiking neurons, action potentials measured at or near the soma are typically thought of as all-or-none signals that invade the full axon terminal to drive synaptic release. Compact, non-spiking neurons, like BCs, are assumed to be nearly isopotential (**Fig. 6**). Thus, somatic voltage is measured as the signal driving synaptic release. There is precedent, however, for both spiking and non-spiking neurons transmitting different signals at different output locations. The degree to which functional divergence occurs in axons has been highlighted as one of the most important questions in neuroscience^43^.

Motor neurons in both rat^44–46^ and spiny lobster^47–49^ propagate spikes down some parts of the axonal arbor but not others in conditions that the authors of these studies argued were physiological. Motor neurons in locusts contain two axonal branches, each with its own axon initial segments that can initiate spikes independently. These spikes often propagate to the opposite branch but can fail to propagate in some conditions^50^. Functional divergence has also been measured in spiking sensory neurons. Leach mechanoreceptors can propagate spikes to different postsynaptic neurons and fail to propagate to others depending on which part of their receptive field is stimulated^51–56^. Auditory afferents in bush cricket can display different frequency tuning in nearby parts of the axonal arbor through a mechanism involving presynaptic inhibition^57^. Mechanisms for the functional compartmentalization of the axonal arbors of spiking neurons have included both intrinsic electrical properties^44–46,58–60^ and the external influence of GABAergic interneurons^61–63^.

In the retina, functional divergence has been demonstrated in several types of non-spiking ACs, including the A17 [ref. ^64^], VGluT3 [ref. ^65^], and starburst^66^. These ACs, however, have substantially larger neuritic arbors than BCs axons, so electrotonic isolation is greater and can better support functional divergence. Some types of BCs in the zebrafish retina have distinct lobular output boutons in different layers of the IPL, which can display different light-driven calcium signals. Differential bouton volume has been suggested as a mechanism for functional divergence in these BCs^67^.

### Possible mechanisms of synapse-specific surround suppression in bipolar cells

Compartmentalization of synaptic release at the micron scale is difficult to reconcile with the canonical view that transmitter release within a neuron is controlled exclusively by presynaptic voltage, especially for a cell type typically modeled as passive and isopotential. Indeed, our modeling suggests that such a passive model of the BC would not support sufficient electrical compartmentalization for much subcellular functional divergence. However, the inclusion of active conductances decreased the membrane resistance to the point that voltage gradients could feasibly contribute to functionally divergent signals within a single BC.

This model contained many assumptions, such as the degree of functionally specific synapse formation and the linearity of the voltage-to-glutamate relationship. If these assumptions are shown to be incorrect, such as functionally indiscriminate BC-RGC synapse formation, the model would overestimate voltage-driven functional divergence. However other inaccuracies, such as a supralinear relation between voltage and glutamate release^41,42^, would cause the model to underestimate voltage-driven functional divergence. Regardless, our model highlights the importance of considering active conductances and challenges the assumption that all output synapses experience the same voltage signal.

Although our modeling suggests the possibility of voltage compartmentalization, chemical compartmentalization within the BC terminal could also contribute to functional divergence. But which molecule(s) could be localized at the micron scale to alter glutamate release? Calcium influx triggers vesicle fusion and glutamate release, and voltage-gated Ca^2+^ channels are clustered near ribbons where they can form Ca^2+^ nanodomains^68^. Strong buffering, including by the release machinery itself, and physical barriers, like the endoplasmic reticulum, the ribbon, and the hundreds of associated vesicles, could limit Ca^2+^ diffusion. Different subtypes of voltage-gated Ca^2+^ channels or associated accessory proteins that modulate their function could be localized to particular synapses, and these barriers to Ca^2+^ diffusion could endow the nanodomains with some degree of independence.

We think the most likely mechanism is that an external, diffusible chemical signal (from ACs to the BC ribbon) causes local regulation of individual synapses. Our pharmacology results showed that GABA_C_ receptors and spiking ACs are required for surround suppression of Pix_ON_ excitation (**Fig. 5b**), but this does not exclude the involvement of another modulator. Perhaps GABA release from spiking ACs causes a moderate hyperpolarization of the BC, which is necessary for any level of surround suppression, but differing levels of surround suppression are achieved through the simultaneous release of an additional modulator which induces a voltage-independent reduction of Ca^2+^ at BC-Pix_ON_ synapses or a voltage-independent enhancement of Ca^2+^ at BC-ON alpha synapses.

Collectively, ACs contain at least 20 different small molecule or peptide transmitters and neuromodulators, and the differential expression of these molecules is one of the primary ways to classify them into different types^69^, yet the functions of most of these molecules in visual processing remain largely unknown. Reports of voltage-independent effects of these substances on calcium levels include activation of Ca^2+^-permeable α7 nicotinic acetylcholine receptors on type 7 BCs^70^, activation of D1 dopamine receptors on rat and mouse BC terminals^71,72^, leading to increased Ca^2+^ levels through PKA-dependent enhancement of L-type Ca^2+^ currents or through PIP2-dependent Ca^2+^ release from internal stores^73^, and regulation of Ca^2+^ current at OFF BC terminals via S-nitrosylation from retrogradely released nitric oxide^74^. While their molecular mechanisms and possible subcellular compartmentalization were not studied, dopamine has been shown to decrease surround suppression in fish BCs^75^, and both agonists and reverse agonists of cannabinoid receptors have been shown to alter the surrounds of mouse ON alpha RGCs^76^.

### Measuring functional divergence at the micron scale

While our interpretation is that functional divergence in BC terminals can occur at the scale of tens of microns, our functional measurements were made at the macroscopic scale of spikes and synaptic currents in RGCs. Functional imaging of calcium or glutamate with genetically encoded indicators could presumably offer more direct measurements at the micron scale. These techniques have been used for studying functional compartmentalization in retinal ACs^64,77,78^, the dendrites of RGCs^79–81^, and even in a recent paper that similarly reported divergence of a different function (direction selectivity) in type 7 BCs^30^.

While these imaging techniques theoretically offer better spatial resolution, they are indirect measures of synaptic function and suffer from their own technical limitations. Calcium imaging revealed functional compartmentalization in A17 ACs where synaptic boutons are separated by ∼20 μm sections of a single, extremely thin (100 nm) neurite^64^. In contrast, a type 6 BC terminal has ∼90 ribbon synapses all within a compact 3D structure mostly lacking anatomical compartmentalization (**Fig. 6a**). In addition to the ever-present issue of the nonlinear relationship between calcium changes and neurotransmitter release, the morphology of these terminals makes measurements of local calcium at the scale of individual synapses with a diffusible indicator infeasible.

Glutamate imaging enables a more direct measurement of the molecule driving postsynaptic current, but it suffers from a different kind of spatial localization problem: uncertainty about the origin of the glutamate. The sensor (iGluSnFR) is present throughout the membrane of each cell in which it is expressed. Thus, it lacks synaptic localization. Expressing iGluSnFR in RGCs could reveal postsynaptic compartmentalization, but it would not reveal whether nearby signals arose from the same or different BCs. Alternatively, expressing iGluSnFR sparsely in the BCs themselves, as was achieved for type 7 BCs via subretinal viral injections^30^, does not guarantee that the measured signals arise from the BCs in which the sensor is expressed given the extremely high density of glutamatergic synapses in the IPL. Of course, any imaging technique in the functioning retina also interferes to some extent with the light responses of the photoreceptors^82^. Laser-induced light exposure is especially problematic when attempting to compare responses to small spots of light within the imaging field (the scale of one or several BCs) to large spots of light that extend beyond the imaging field.

Instead of functional imaging, we used electrophysiology to ascertain the presynaptic origin of the divergence in surround suppression between Pix_ON_ and ON alpha RGCs (**Figs. 1-3 & Supplementary Figs. 2-5**). We then used SBFEM and confocal imaging to determine that these RGCs share input from the same set of BCs (**Fig. 4 & Supplementary Fig. 7**). Thus, while we did not directly measure different functional signals within the terminals of a single BC, we showed that different glutamate release profiles are experienced by RGCs that collect from the same population of BCs.

### Implications for visual processing in bipolar cells

A decade ago, RGC spike recordings during current injections into single salamander BCs suggested that individual BCs could, at least indirectly, transmit different functional signals to different RGCs; however, it remained unclear to what extent postsynaptic mechanisms, ACs, or gap junctions were involved^83^. These authors and others^84^ have speculated about the vast computational power of a neural network in which individual connections between neurons could have some degree of functional independence. Our results demonstrate that, indeed, one of the most canonical retinal computations, surround suppression, can manifest within a neuron whose output synapses are on average less than 25 μm apart.

We focused on type 6 BCs (**Fig. 5e, Fig. 6**, and **Supplementary Fig. 7**), but type 7 BCs also provide a substantial input to Pix_ON_ and ON alpha RGCs (**Fig. 4h-j**), and functional divergence of direction selectivity in their terminals has been measured by glutamate release^30^. Rather than an exception, functional divergence may be the rule in mouse (and perhaps other mammalian) BCs. Importantly, one cannot necessarily measure functional divergence with a single stimulus paradigm^30^. Since the difference we measured was in the degree of surround suppression, we would not have measured it with spatially uniform stimuli or when analyzing only a single spot size at a time. This could help explain the lack of evidence for subcellular processing in a previous study of mouse BC glutamate release^19^, though the same researchers did find evidence for functional divergence in BCs with improved analysis methods^85^.

Functional specialization at the subcellular scale is noteworthy in the context of interpretations of ultrastructural (connectomics) datasets, where the mouse retina has been a model for linking circuit structure to function^7,86,87^. EM reconstructions allowed us to quantify BC inputs to Pix_ON_ and ON alpha RGCs (**Fig. 4**) and to measure details of the locations of synapses (**Figs. 5f, 6a**), but the main conclusion of our study suggests that one should be cautious in interpreting similar patterns of synaptic connectivity as a proxy for similar function.

## Methods

### Ex vivo retina preparation

Mice of either sex aged 6 - 36 weeks were used for recordings and imaging. For experiments requiring labeled type 6 BCs (**Figs. 4-5** and **Supplementary Fig. 7**), CCK-ires-Cre/Ai14 mice were used (Jackson Lab Strain # 012706 / 007914). All other experiments used wild-type mice (C57BL/6, Jackson Lab Strain # 000664).

Whole mount retinas were prepared in a similar manner to previous publications^8,88–93^. In short, dark-adapted mice were sacrificed, and retinas were dissected under infrared illumination (940 nm). The intact retina was flat-mounted photoreceptor side down on a poly-D-lysine-coated glass coverslip and placed in a recording chamber. Retinas were perfused with oxygenated Ames medium at 32°C at a rate of 10 mL/min throughout the experiment. Animals were sacrificed following animal protocols approved by the Center for Comparative Medicine at Northwestern University.

### Visual stimulation

Visual stimuli were generated with a 912 × 1140 pixel DLP projector (1.3 μm/pixel) at a 60 Hz frame rate using a blue LED (450 nm) focused on the photoreceptor outer segments. Light intensities are reported in rhodopsin isomerizations per rod per second (R*/rod/s). Visual stimuli had intensity values of 200-300 R*/rod/s and background intensity values of ∼0.3 R*/rod/s unless otherwise noted (**Supplementary Figs. 2** and **5**). Each cell’s receptive field center was determined by flashing horizontal and vertical bars at different locations, and all subsequent stimuli were centered on the location that elicited maximal responses. Surround suppression was probed using a pseudorandom sequence of 12 spot sizes (diameters logarithmically spaced from 30-1200 μm) each presented for 1 second.

### Cell-attached and whole-cell recordings

All recordings were obtained using a 2-channel patch-clamp amplifier (Multiclamp 700B, Molecular Devices) sampling at 10 kHz. Spike trains were recorded using glass pipettes (2–3MΩ) filled with AMES solution in cell-attached configuration. Voltage-clamp recordings were performed using glass pipettes (4– 6MΩ) filled with a cesium-based intracellular solution (105 mM Cs methanesulfonate, 10 mM TEA-Cl, 20 mM HEPES, 10 mM EGTA, 2 mM QX-314, 5 mM Mg-ATP, and 0.5 mM Tris-GTP; ∼277 mOsm; pH ∼7.32 with CsOH). Voltage was corrected for the liquid junction potential (−8.6 mV) and the cell was clamped to the reversal potential of chloride (−60 mV) to measure excitatory conductances or the reversal potential of glutamate-induced cation currents (+20 mV) to measure inhibitory conductances. Current clamp recordings and cell fills of neurobiotin were performed using glass pipettes (4–6MΩ) filled with a potassium-based intracellular solution (125 mM K-aspartate, 10 mM KCl, 1 mM MgCl2, 10 mM HEPES, 1 mM CaCl2, 2 mM EGTA, 4 mM Mg-ATP and 0.5 mM Tris-GTP; 77 mOsm; pH ∼7.15 with KOH).

### Dynamic clamp recordings

Dynamic clamp hardware and software were implemented as described in Desai et al. (2017). Pix_ON_ and ON alpha excitatory and inhibitory conductances were recorded in response to 200, 600, and 1200 μm diameter spots of light. New RGCs were then patched in whole-cell current-clamp configuration, the previously recorded conductances were simulated via current injections, and the resulting spike train was recorded. In each experiment, excitatory and inhibitory conductances were simultaneously simulated, and these excitatory and inhibitory conductances were equally scaled for each cell to best reproduce the cell’s response to a real preferred spot visual stimuli.

For **Figure 2b-g**, the paired excitatory and inhibitory conductances were always derived from the same size spot stimuli (e.g., if simulating excitation evoked by a 200μm spot, inhibition evoked by a 200 μm spot was simultaneously simulated). The “Preferred size” response was taken as the maximal spiking response observed during the simulation of all excitatory and inhibitory conductance pairs (200 μm, 600 μm, and 1200 μm). “Full-field” responses were taken as the spiking response when simulating excitation and inhibition recorded during 1200 μm spot stimuli.

For **Figure 2h,i**, “Exc_pref_” and “Inh_pref_” refer to the Pix_ON_ excitatory and inhibitory conductances that were found to elicit the maximal spiking response in **Figure 2b,d**. Conversely, “Exc_ff_” and “Inh_ff_” refer to the Pix_ON_ excitatory and inhibitory conductances recorded during the presentation of a 1200 μm diameter light spot.

### Pharmacology

Intrinsic light responses were measured by providing full-field light stimuli while voltage clamping at -60 mV during bath application of L-AP4, DNQX, and D-AP5 to block photoreceptor-driven light response^94^. See **Supplementary Table 1** for a complete listing of pharmacological agents and their targets.

### Physiology analysis

RGC spiking responses were measured as the average spike rate during the 1-second light stimulus. RGC conductance responses were measured as the total charge transfer during the 1-second light stimulus. The preferred size response (R_preferred size_) was defined as the maximal response measured during the presentation of all sizes of spot stimuli (30 - 1200 μm diameter). The full-field response (R_full-field_) was defined as the response recorded during the presentation of the largest stimulus spot (1200 μm diameter). Suppression was calculated as

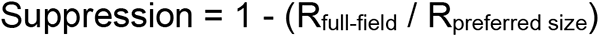

### Two-photon imaging

RGCs were filled with AlexaFluor 488 via a whole-cell patch pipette. Images were collected through a ×60 water immersion objective (Olympus LUMPLan FLN 60x/1.00 NA) using 980 nm two-photon laser excitation (MaiTai HP, SpectraPhysics).

### Immunohistochemistry

Retinas were fixed at room temperature for 15 min in 3% paraformaldehyde (Electron Microscopy Sciences) and then blocked at room temperature for 2 hours in 3% Normal Donkey Serum (Jackson Labs) and 0.5% Triton (Sigma) in Phosphate Buffer.

Retinas were then incubated with primary antibodies for 5 days at 4°C. After washing, retinas were incubated with secondary antibodies for 2 days at 4°C. All antibodies were diluted 1:500. Retinas were then mounted on glass coverslips using Vectashield Antifade (Vector Labs).

### Confocal imaging

RGCs were filled with Neurobiotin tracer (Vector Laboratories, SP-1150, ∼3% w/v potassium-based internal solution) and fixed in 3% paraformaldehyde solution. After performing immunohistochemical labeling, tissues were imaged on a Nikon A1R laser scanning confocal microscope through a ×40 or ×100 oil immersion objective (Nikon Plan Apo VC ×40/×60/1.4 NA).

### Quantification of RGC morphology

From both two-photon and confocal images, soma diameter was calculated by tracing an outline of the soma using ‘Freehand Selections’ and solving for diameter in FIJI. Similarly, the dendritic diameter was measured by drawing a convex polygon around the tips of the dendrites in a flattened view of the image. Average branch length, number of branches, and total dendritic length were calculated by tracing the RGC dendrites using the SNT plug-in in FIJI and its built-in analysis tools^95^.

Stratification analysis was performed by measuring dendrite depth in the IPL in relation to the immunohistochemically labeled ChAT bands (Starburst amacrine dendrites). Custom MATLAB software (Nath and Schwartz, 2016) based on a published algorithm ^96^ were used to flatten the image prior to analysis.

M5 and M4 RGC morphological data were provided by Professor David Berson and are published in Stabio, et al.^94^ and Estevez et al.^17^, respectively.

### Correlative Fluorescence and serial block-face scanning electron microscopy (SBEM)

Neighboring Pix_ON_ and ON Alpha RGCs with overlapping dendritic arbors were physiologically identified in a mouse line with fluorescently labeled type 6 BCs (CCK-ires-Cre/Ai14^22,23^). After verifying that the two ganglion cells had differing levels of surround suppression in their spiking response, both ganglion cells were filled with Alexa 488. Two-photon volume images of the RGCs overlapping dendrites and the type 6 BC axonal arbors were then acquired. The retina was fixed with 1.5% glutaraldehyde and 2.5% paraformaldehyde in 0.1M Na Cacodylate buffer for 10 minutes. The retina was washed with 0.1M Na Cacodylate buffer and transferred to 4% glutaraldehyde for 4 hours at 4° C to stiffen the tissue.

We utilized the previously published near-infrared branding technique ^97^ to burn fiducial markers into the retina with the two-photon laser (860 nm, ∼100 mW), allowing for the alignment of two-photon images with electron microscopy volumes. The tissue was prepared for SBEM according to the protocol described previously ^98^. Image stacks were acquired using a VolumeScope SEM (Apreo, Thermo Fisher Scientific) at a voxel size of 5 × 5 × 50 nm^3^.

### Volume reconstruction and Image analysis

EM image stacks were aligned and registered using ImageJ/TrakEM2^99^. The neuronal processes were traced and segmented using AreaTree or AreaList function, whereas synapses were segmented using AreaList function in TrakEM2. The 3D objects of either traced skeletons or surface segmentations were visualized in either 3D view in ImageJ or exported to and rendered in Amira (Thermo Fisher Scientific).

The somata of both Pix_ON_ and ON alpha RGCs were located according to the fiducial markers. We traced the dendritic arbors of both RGCs within the limit of the EM volume. BC synapses were identified by the presence of a presynaptic ribbon apposed to the postsynaptic dendrites of both RGCs. Presynaptic amacrine cell contacts were identified by the presence of clusters of synaptic vesicles apposed to BC axonal terminals. Presynaptic BCs were reconstructed, and their type was determined according to their stereotyped morphology^100–102^. Type 6 BCs were further confirmed by the presence of the fluorescence marker in the corresponding 2-photon volume.

We identified all the presynaptic (amacrine) inhibitory sites on the BC bouton where the ribbon synapses resided. For each ribbon, the Euclidean distances between the ribbon and all the presynaptic inhibitory sites were measured, and the inhibitory synapse with the shortest distance was identified.

### Bipolar cell summation over RGC dendrites model

We modeled RGC excitation across spot sizes as the summation of BC subunits sampled across the RGC’s dendritic arbor. To do this, a skeleton of the RGCs dendritic arbor was provided to the model, and excitatory input synapses were randomly assigned along the length of the dendritic skeleton (0.3 μm / synapse^103,104^). At each of these synapses, a BC was assigned, and its receptive field was centered on that synapse.

RGC excitatory responses across spot sizes were predicted by presenting virtual spots of multiple sizes centered at the centroid of the ganglion cell dendritic field and calculating each BC’s activation as the overlap of its receptive field with the presented spot. RGC excitatory conductances were then modeled as the linear sum of each BC’s activation. Both experimentally measured and model-predicted excitatory responses were normalized across spot sizes by the maximal response.

The BC receptive field was modeled as a circular difference of gaussians^105^ with 3 parameters; center size (σ_c_), surround size (σ_s_), and center-to-surround ratio (CSR). While σ_c_ was fixed at 22 μm [ref. ^104^], σ_s_ and CSR were obtained by minimizing the mean absolute error between the model output and the experimentally recorded RGC excitatory responses across all spot sizes. Error was minimized using the Interior-point optimization algorithm, and initial values of 100 μm for σ_s_ and 1 for CSR. 6 RGCs were simultaneously fit for each estimation of σ_s_ and CSR. Cross-validation was performed on the remaining RGCs. When fitting to both Pix_ON_ and ON alpha RGCs, 3 Pix_ON_ and ON alpha RGCs were used for fitting. Four hundred random fitting combinations of the 14 Pix_ON_ and 8 ON alpha RGCs were performed to obtain average cross-validation values.

The model was written using MATLAB 2022a. Code and data are available at https://github.com/davidswygart/rgc_bipolar_dog.

### NEURON compartment model of type 6 bipolar cell

Modeling was performed using Python 3.8 and NEURON 8.0 [ref. ^28^]. SBSEM reconstructions were imported to NEURON using NEURON’s *Import3d* tool. To simulate light-evoked activation, the axon terminals were depolarized to ∼-35 mV by activating synapses on its dendrites (0 mV reversal potential^41^). Each inhibitory synapse (reversal potential = -60 mV) was then activated with enough conductance to hyperpolarize the membrane to -45 mV the location of the activated inhibitory synapse^41,106^. When measuring the effect of a single inhibitory synapse (**Fig. 6c** & **7c**), hyperpolarization was measured for all 91 ribbon output synapses in response to the activation of each of the 120 inhibitory synapses (a new simulation was performed for each inhibitory synapse). When testing the simultaneous activation of inhibitory synapses (**Fig. 6d-f** & **Fig. 7d**), the same 120 inhibitory synapses were sequentially activated, but the additional N number of nearest inhibitory synapses (by path distance) were also simultaneously activated.

Inhibition-induced hyperpolarization was measured for each ribbon as the change in membrane potential caused by activation of the inhibitory synapses. All measurements were acquired after the membrane had reached a steady-state (200 ms). To calculate Q4→Q1 inh. decay, ribbons were sorted by the magnitude of their hyperpolarization. The 91 ribbons were then split into quartiles (23 ribbons in each quartile), and the average hyperpolarization was calculated for the top (Q4) and bottom (Q1) quartiles. Inhibitory decay was then calculated as 100% - Q1 / Q4.

Key model parameters can be found in **Table 3**. Model stability was tested by measuring inhibitory voltage decay across a range of key model parameters (**Supplementary Fig. 9**). Whenever model parameters were altered, excitatory and inhibitory conductances were adjusted to maintain the same membrane potentials at the inhibitory synapse (−35 mV → -45 mV).

To better enable comparisons of our model to previous literature, we fit length constants for each inhibitory synapse (**Supplementary Fig. 9**). To do this, hyperpolarization was measured at the activated inhibitory synapse (∆V_inh_) and at each ribbon output synapse (∆V_rib_). The length constant (λ) for each inhibitory synapse was estimated by fitting

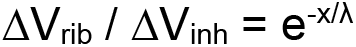

where x was the path distance between each ribbon and the activated inhibitory synapse.

**Supplementary Fig. 9** shows that the median length constant of the inhibitory synapses in our passive BC model was 616 ± 138 μm (± median absolute deviation), which is similar to values estimated in previously published passive models of rod BCs^29^ and type 7 ON cone BCs^30^. In our active BC model that contained L-type Ca^2+^ channels^36,37^, K_V+_ channels^38^, and HCN_2_ channels^39,40^, the median length constant of the inhibitory synapses was 82 ± 13 μm (median ± median absolute deviation).

Code and data are available at,https://github.com/davidswygart/T6_NEURON_python

### Statistical tests

Statistics and data representation are reported in figure legends. In short, data are reported as mean ± SEM. Differing means were assessed with Welch’s t-test for unpaired data, paired two-sample Student’s t-test for paired data, and two-way ANOVA for multivariate data. Comparisons of proportions were assessed with a two-proportions z-test with Holm-Bonferroni correction. Differing continuous distributions were assessed with Kolmogorov–Smirnov tests.

## Acknowledgments

We are thankful to all Schwartz lab members for their feedback and technical assistance throughout the project. We would like to thank Tiffany Schmidt and Anna Vlasits for their feedback and comments on the manuscript and David Berson for sharing M4 and M5 morphological data^17,94^. Funding for this research was provided by National Institutes of Health grants 5F31EY030344-02 and 5R01EY031029-02. Additionally, S. Takeuchi was supported by the Graduate Research Abroad in Science Program (GRASP) of The University of Tokyo and the Graduate Program for Leaders in Life Innovation (GPLLI).

## Author Contributions

D.S. and G.W.S. designed the experiments. D.S. performed experiments and analyzed results related to electrophysiology, flourescent imaging, and mathematical modeling. W-Q.Y, S.T., R.O.W. performed experiments and analyzed results related to Serial Blockface Electron Microscopy. D.S. and G.W.S wrote the manuscript with feedback from W-Q.Y, and R.O.W.

