## Extended Data Figures for "A presynaptic source drives differing levels of surround suppression in two mouse retinal ganglion cell types"

### Supplementary Figures

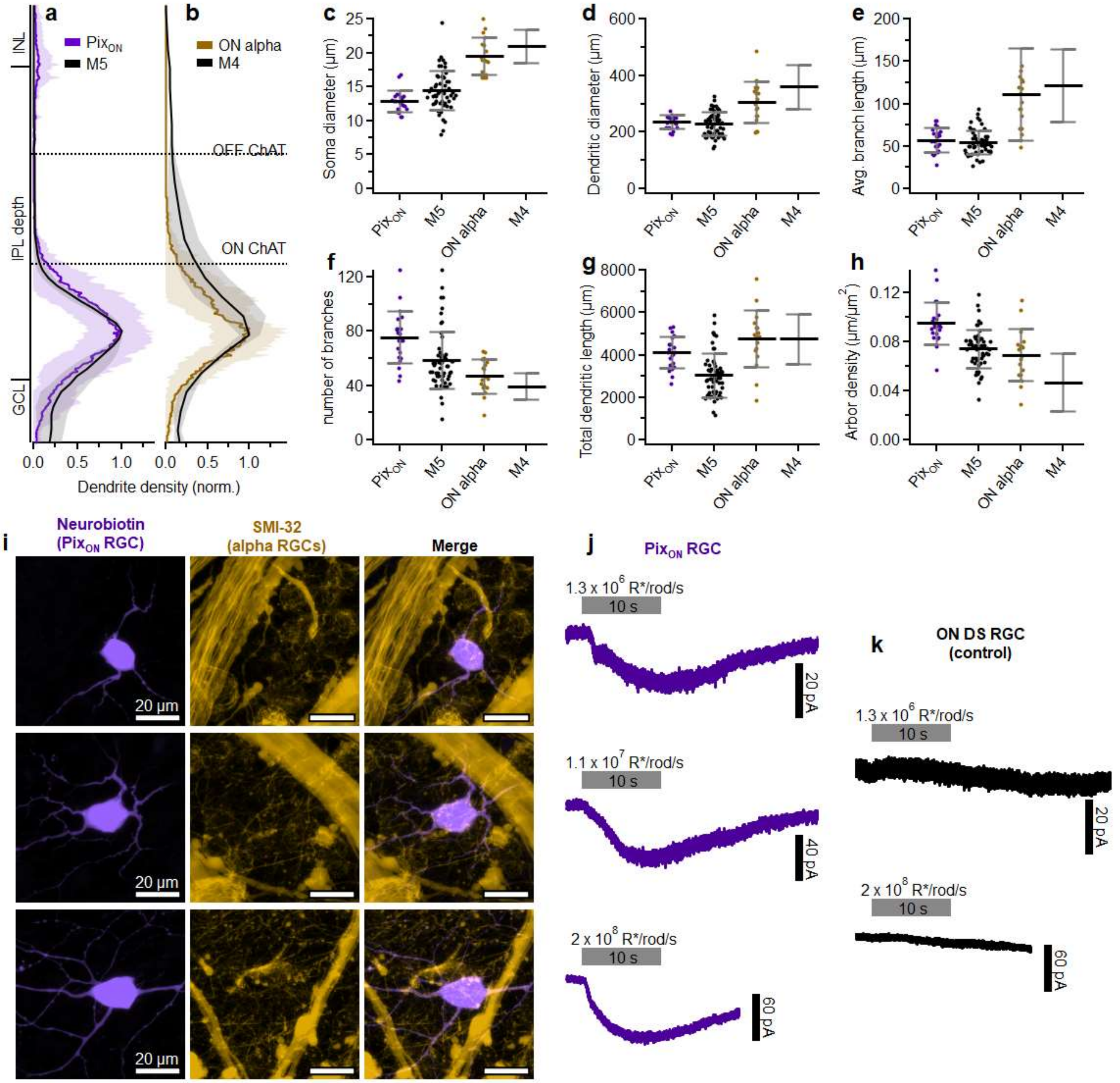


**Fig. 1 | Pix$_{ON}$ and ON alpha RGCs exhibit unique morphology and correspond to M5 and M4 RGCs.**
**a**, Dendritic stratification of Pix$_{ON}$ (n=19) and M5 (n=2) RGCs within the inner nuclear layer (INL), inner plexiform layer (IPL), and ganglion cell layer (GCL). Dotted lines refer to the ON and OFF choline acetyltransferase (ChAT) bands used to determine stratification. Shading and error bars indicate ± standard deviation. **b**, Same as **a**, but comparing ON alpha (n=10) and M4 (n=2) dendritic stratification. **c**-**h**, Comparison of soma diameter (**c**), dendritic diameter (**d**), average dendritic branch length (**e**), total number of dendritic branches (**f**), total dendritic length (**g**), and arbor density (**h**) between Pix$_{ON}$ (n=22), M5 (n=56), ON alpha (n=18), and M4 (n=27) RGC types. Arbor density (**h**) was calculated as the total dendritic length normalized by dendritic area. Dots indicate data from individual cells. Bar plots indicate average ± std. **a**-**h**, M5 and M4 RGC morphological data were provided by Professor David Berson and are published in Stabio, et al. (2018) and Estevez et al. (2012), respectively. **i**, *En-face* view of three different Pix$_{ON}$ RGC somas visualized by neurobiotin fill (left), SMI-32 staining to mark alpha RGCs (middle), and merged images. **j**, Intrinsic photocurrents measured by voltage-clamp recordings ($V_{CMD}$ = -60 mV) during pharmacological blockade of retinal synapses (L-AP4, DNQX, and D-AP5). Gray bars indicate 10-second full-field light step. Light intensity is reported in rhodopsin isomerizations per rod per second (R*/rod/s). Currents were measured from the same cells as in **i**. **k**, Same as **j**, but recorded from ON direction-selective RGCs, which are not expected to exhibit intrinsic photocurrents.

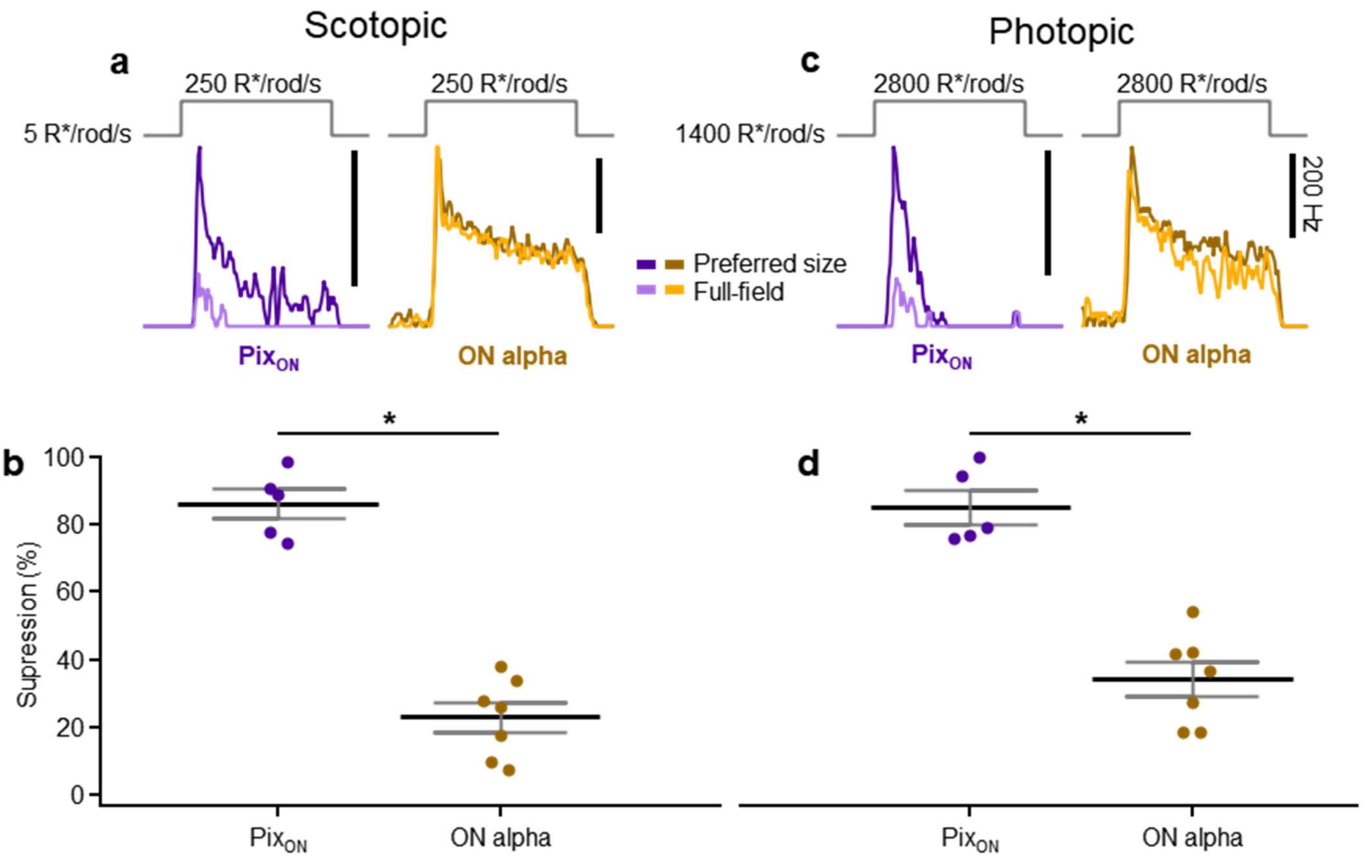


**Supplementary Fig. 2 | Pix$_{ON}$ and ON alpha RGCs have differing levels of surround suppression in both scotopic and photopic conditions.**

**a**, Example Pix$_{ON}$ (purple) and ON alpha (brown) peristimulus time histograms to preferred size and full-field light spot stimuli. The stimulus occurred within the scotopic luminance regime, stepping to a light intensity of 250 rhodopsin isomerizations per rod per second (R*/rod/s) from a background intensity of ~0.3 R*/rod/s for 1 second. **b**, Surround suppression in Pix$_{ON}$ (n=5) and ON alpha (n=7) RGCs to scotopic stimuli. Dots indicate data from individual cells. Bar plots indicate average ± s.e.m., *P<0.05, paired two-sample Student's *t*-test. **c**, Same as **a**, but the stimulus occurred within the photopic luminance regime, stepping from 1400 to 2800 R*/rod/s for 1 second. **d**, Surround suppression to photopic stimuli for the same cells as in **b**.

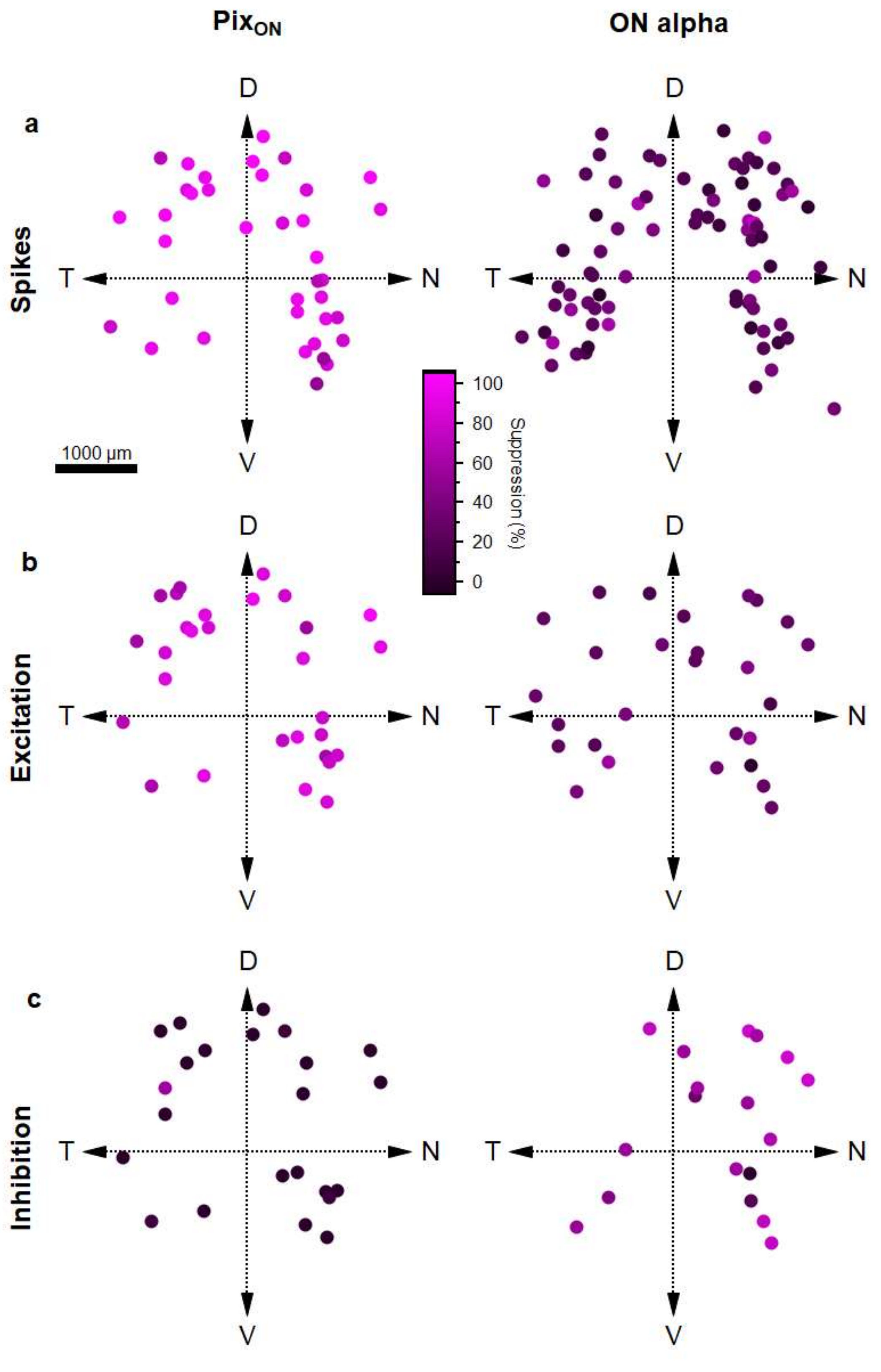


**Supplementary Fig. 3 |Pix$_{ON}$ RGCs have stronger surround suppression than ON alpha RGCs across retinal locations.** **a**, Surround suppression of Pix$_{ON}$ (left) and ON alpha (right) spiking responses plotted by retinal location. Dots indicate the location of individual cells plotted on a dorsal (D) / ventral (V) / temporal (T) / nasal (N) coordinate scheme of the retina. Pix$_{ON}$ (n=38), ON alpha (n=79). **b**, Same as **a**, but for surround suppression of excitatory conductances. Pix$_{ON}$ (n=30), ON alpha (n=27). **c**, Same as **a**, but for surround suppression of inhibitory conductances. Pix$_{ON}$ (n=24), ON alpha (n=18).

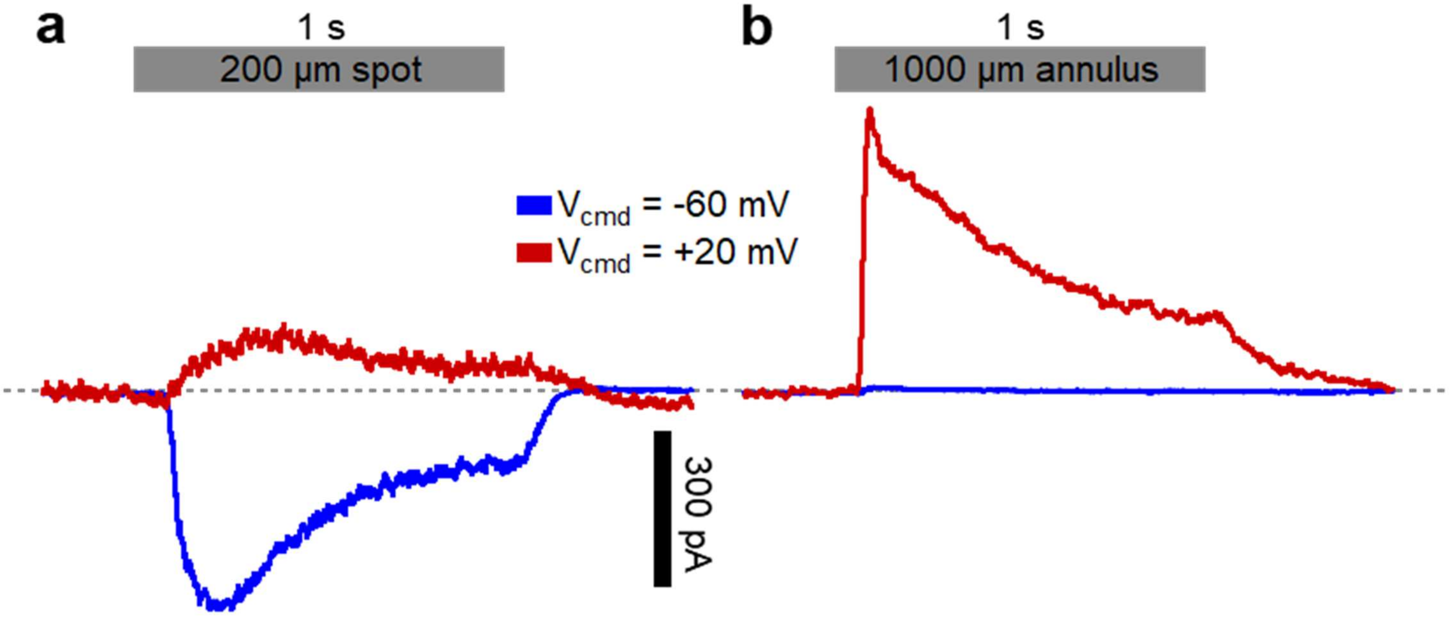


**Supplementary Fig. 4 |Verification of voltage-clamp isolation of excitatory and inhibitory currents in a Pix$_{ON}$ RGC.** **a**, Synaptic currents evoked by a 1-second light step of a 200 µm diameter spot while voltage clamping at -60 mV (blue) or +20 mV (red). This stimulus primarily activated the RGC's receptive-field center, which has strong excitatory input and weak inhibitory input. **b**, Same as **a**, but the visual stimulus was an annulus with an inner diameter of 1000 µm and an outer diameter of 1200 µm. This stimulus primarily activated the RGC's receptive-field surround, which has weak excitatory input and strong inhibitory input.

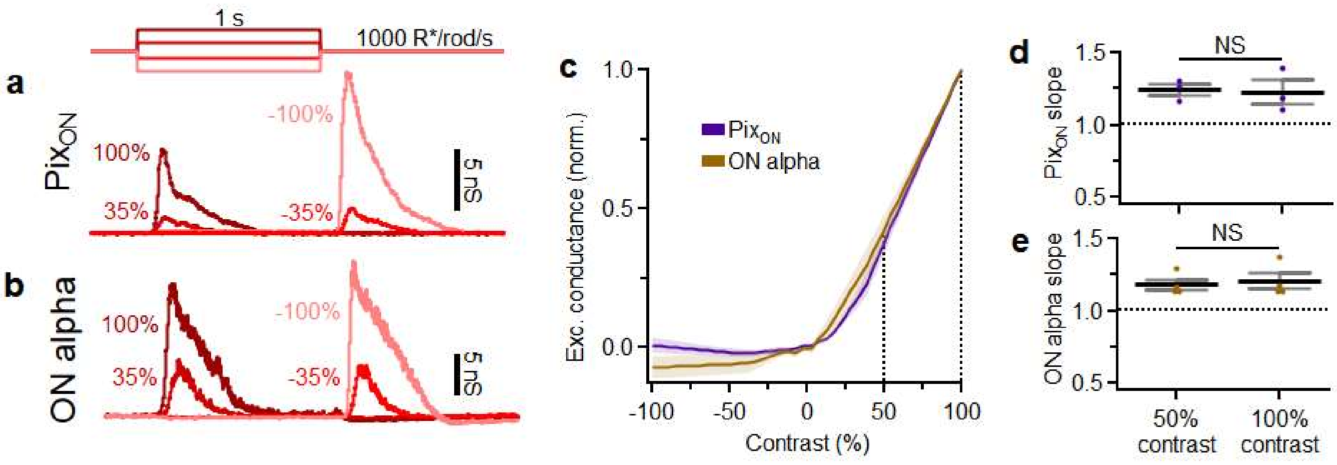


**Supplementary Fig. 5 | Pix$_{ON}$ and ON alpha RGCs exhibit similar contrast response functions in their excitatory conductances.**
**a**, Example Pix$_{ON}$ excitatory conductances evoked by stimulation of positive and negative contrast steps from a background illumination of 1000 rhodopsin isomerizations per rod per second (R*/rod/s). **b**, Same as **a**, but recorded from an ON alpha RGC. **c**, Excitatory responses measured across a range of contrast steps for Pix$_{ON}$ (n=3) and ON alpha (n=3) RGCs. **d**, Equal slopes of the Pix$_{ON}$ contrast response (from **c**) indicate that the Pix$_{ON}$ excitatory responses have not begun saturating at 100% contrast compared to 50% contrast. **e**, Same as **d**, but for ON alpha RGCs. Dots indicate data from individual cells. Bar plots indicate average ± s.e.m., NS P>0.05, paired two-sample Student's *t*-test.

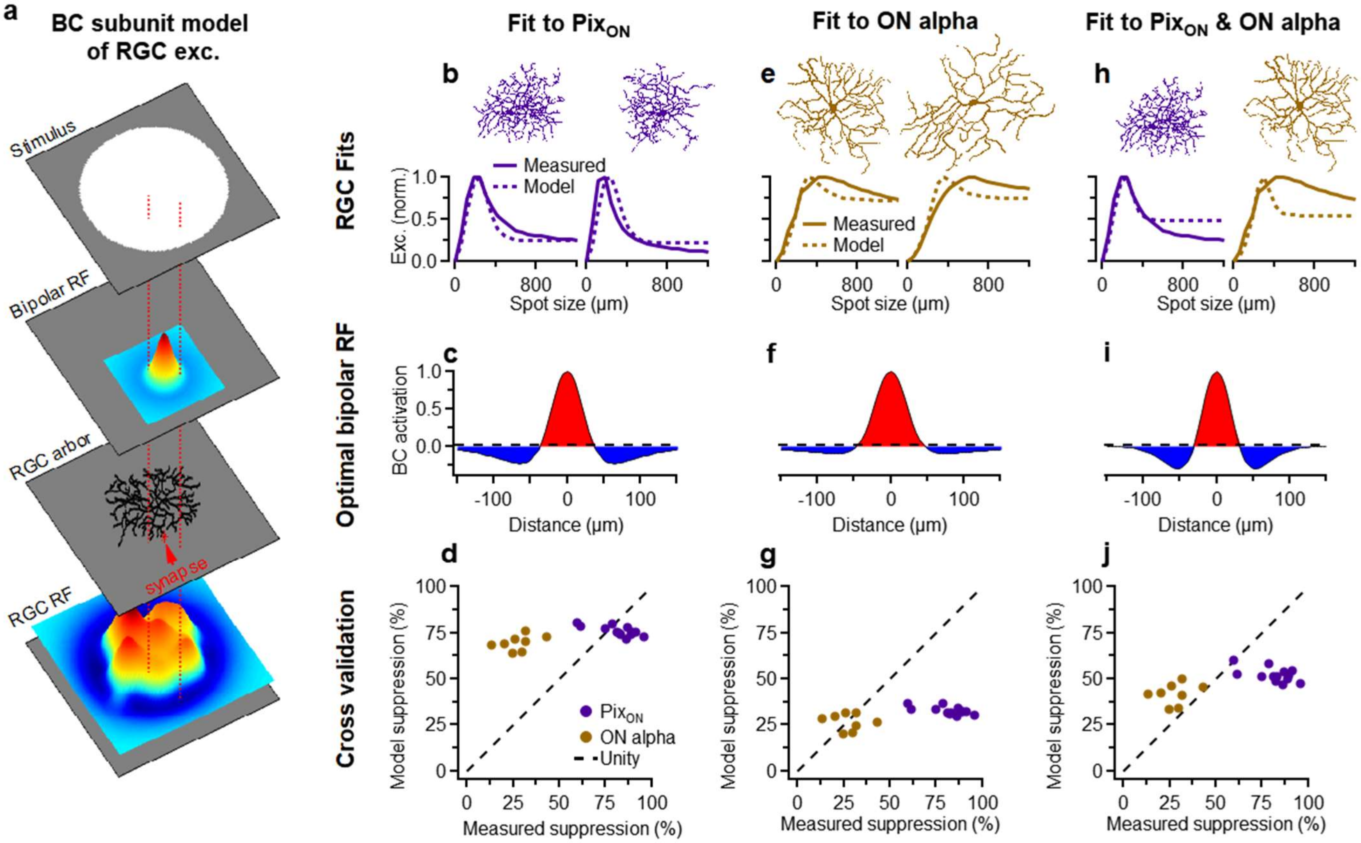


**Supplementary Fig. 6 | A bipolar subunit model of RGC excitation suggests differing bipolar receptive fields are necessary to evoke the differing level of surround suppression observed.**
**a**, Schematic illustrating the BC subunit model of RGC excitation. The RGC receptive field (RGC RF) is constructed from BC receptive fields (Bipolar RF) randomly sampled across its dendritic arbor. RGC excitation is modeled as the overlap of the RGC receptive field with a virtual stimulus. **b**, Two example Pix$_{ON}$ dendritic arbors (*top*) and their corresponding excitatory conductances (*bottom*, solid line) used to fit the BC RF in **c**. Fitting was performed simultaneously on 6 Pix$_{ON}$ RGCs. Dotted lines indicate the model-predicted excitatory responses across spot sizes when using the BC RF in **c**. **c**, The BC receptive field that minimized the absolute error between measured and model-predicted excitatory responses from **a** (see methods for details). **d**, Experimentally measured surround suppression from Pix$_{ON}$ (n=14) and ON alpha (n=8) RGCs plotted against the average surround suppression predicted by the model when cross-validating against Pix$_{ON}$ and ON alpha RGCs not used for fitting the BC RF. Note: alignment to unity indicates perfectly accurate model prediction. **e-g**, same as **b-d**, but fitting to 6 ON alpha RGCs. **h-j**, same as **b-d**, but simultaneously fitting to 3 Pix$_{ON}$ RGCs and 3 ON alpha RGCs.

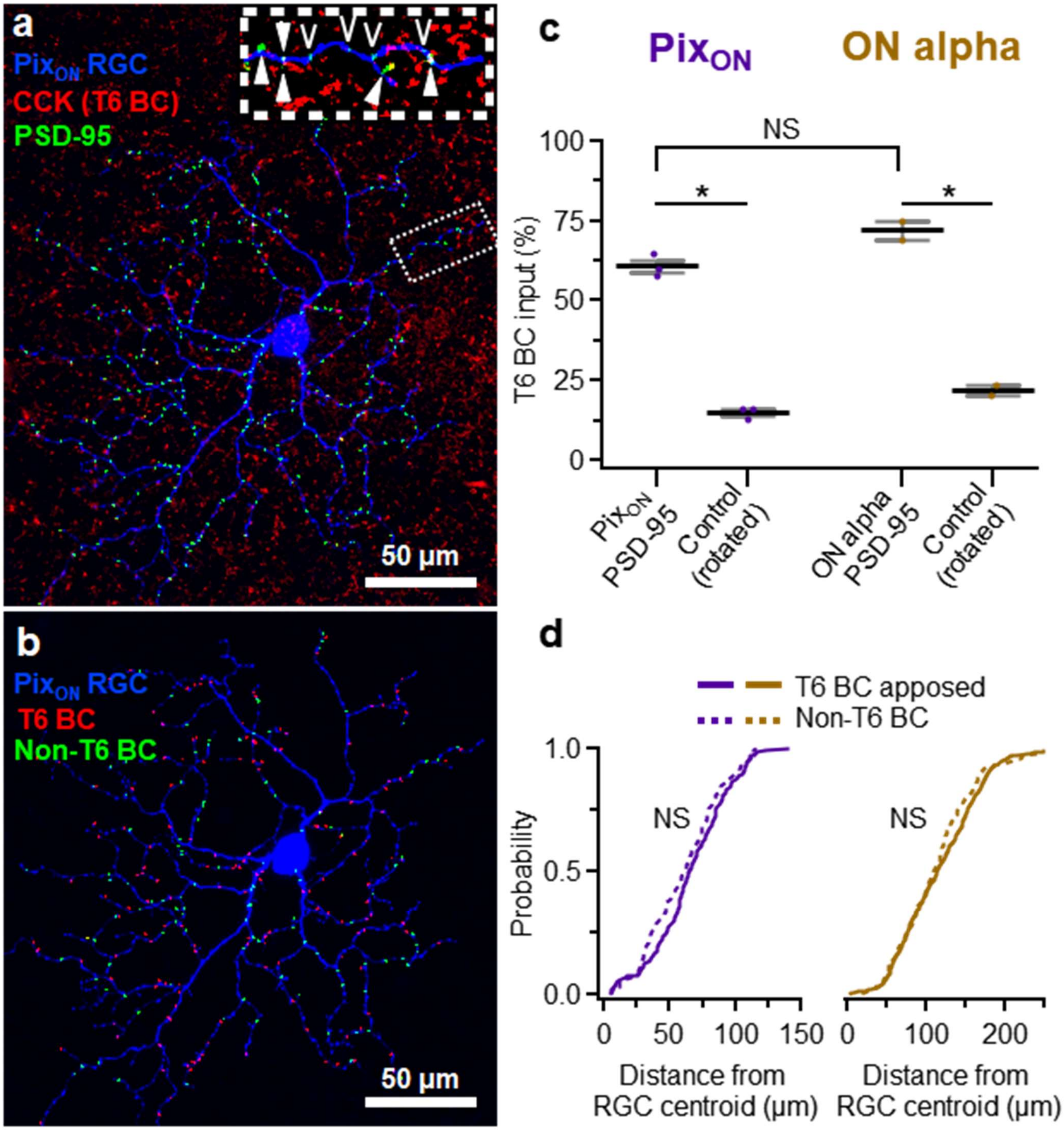


**Supplementary Fig. 7 | PSD95 puncta apposed to type 6 bipolar cell terminals throughout the dendritic arbors of Pix$_{ON}$ and ON alpha RGCs.**

**a**, *En-face* view of a neurobiotin-filled Pix$_{ON}$ RGC (blue) in the CCK-ires-Cre/Ai14 mouse line which labels type 6 BCs (T6 BC, red). Postsynaptic density protein 95 (PSD95, green) is immunohistochemically labeled to identify excitatory synapses on the RGC dendrite. Inset shows a zoomed-in view of the Pix$_{ON}$ dendrite in which some PSD95 puncta are apposed to a T6 BC axon terminal (white closed arrow), while other PSD95 puncta are not apposed to a T6 BC axon terminal (white open arrow). **b**, Same Pix$_{ON}$ RGC as in **a**, but all PSD95 puncta have been identified as apposed (red) or not-apposed (green) to a T6 BC terminal. **c**, Percentage of PSD95 puncta apposed to a T6 BC within the Pix$_{ON}$ (n=3) and ON alpha (n=2) RGCs dendritic arbor. To estimate the chance probability of PSD95 puncta overlapping with T6 BC terminals, we performed a control analysis in which the PSD95 puncta image channel was rotated 90° compared to the T6 BC image channel. *$P<0.05$, Welch's t-test was used for comparison of Pix$_{ON}$ to ON alpha. Paired two-sample Student's *t*-test was used to compare the experimental group to a rotated control. **d**, Cumulative probability of distances between PSD95 puncta and the centroid of the RGC dendritic arbor, plotted for both T6 BC apposed and non-T6 BC apposed PSD95 puncta. Differing distributions of these distances would indicate that the proportion of T6 BC apposed and non-T6 BC apposed PSD95 vary by dendritic eccentricity. These distributions were not found to differ significantly for either Pix$_{ON}$ (T6 n=163, non-T6 n=110) or ON alpha RGCs (T6 n=461, non-T6 n=157). $P>.05$, Kolmogorov–Smirnov test.

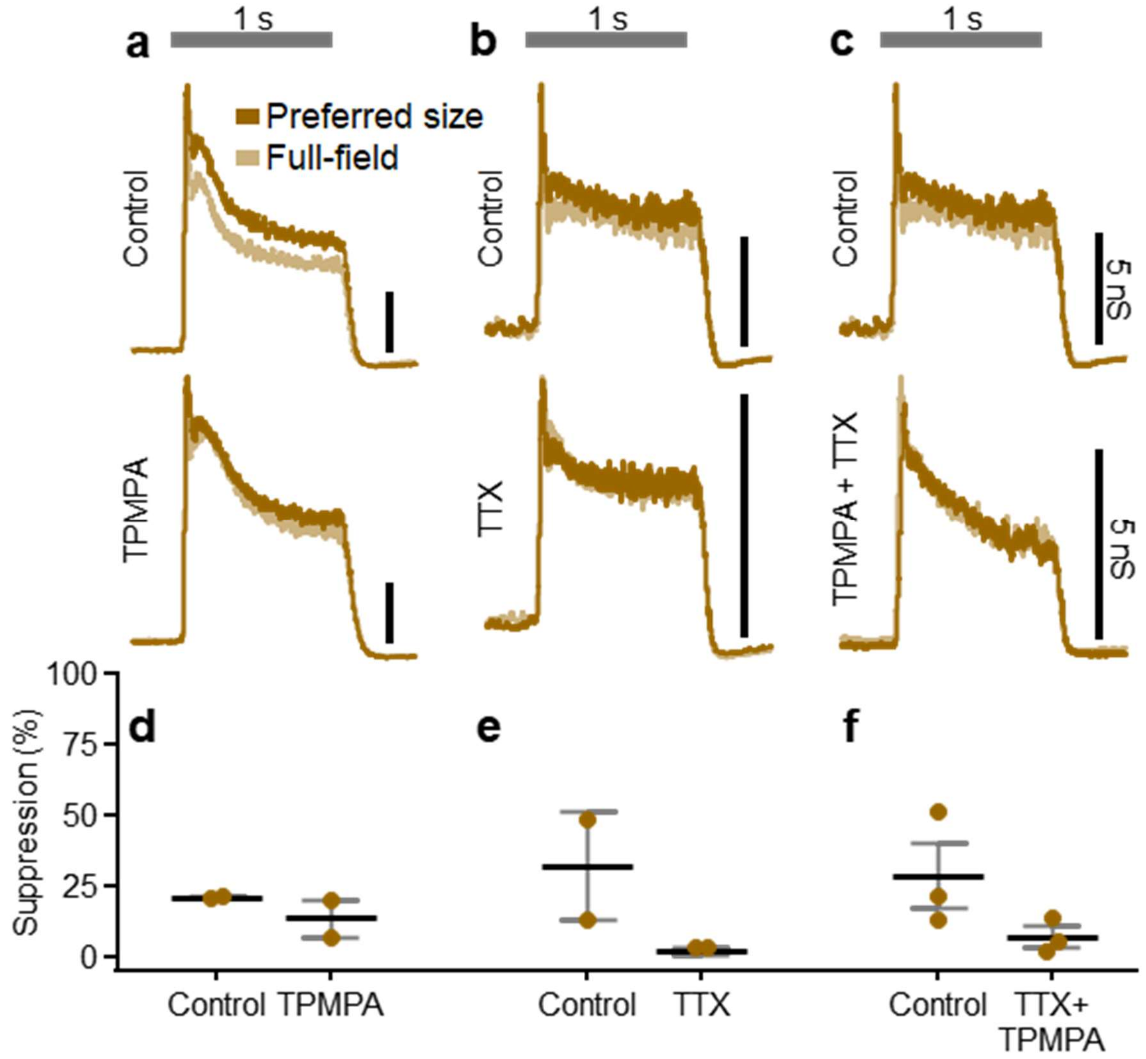


**Supplementary Fig. 8 | ON alpha surround suppression of excitation remains weak in pharmacological block of GABA$_C$ receptors and Na$_V$ channels.**

**a**, ON alpha excitatory conductances evoked before (top) and after (bottom) bath application of the GABA$_C$ receptors antagonist TPMPA. The gray horizontal bar indicates a 1-second presentation of either the preferred size (dark brown) or full-field (light brown) spot stimuli. **b**, Same as **a**, but during bath application of the Na$_V$ channel blocker TTX. **c**, Same as **a,** but during dual application of TPMPA and TTX. **d**, Surround suppression of ON alpha excitatory conductances in control conditions and during bath application of TPMPA (n=3). **e**, Same as **d**, but during bath application of TTX (n=2). **f**, Same as **d**, but during dual application of TPMPA and TTX (n=3). **d-f**, Dots indicate data from individual cells. Bar plots indicate average ± s.e.m.

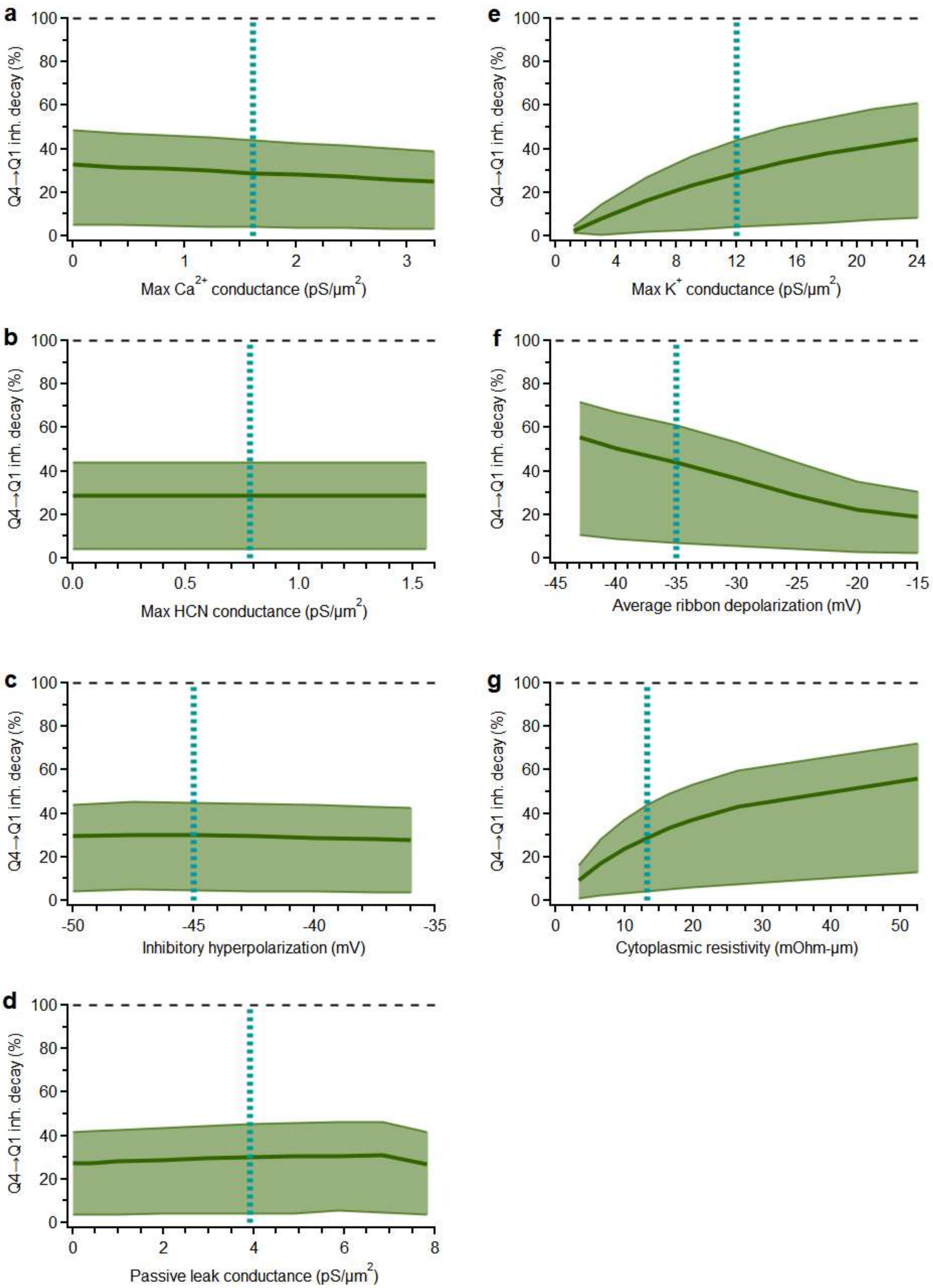


**Supplementary Fig. 9 | Cable model consistency over a range of parameter values.**
**a-d**, Percent decrease of hyperpolarization from the top to the bottom quartile of ribbons when activating a single inhibitory synapse plotted against a range of parameter values (see **Fig. 6**). Thick lines indicate the median decay across all sets of inhibitory synapses activated. Shading indicates the range (maximum to minimum) of inhibitory decay across all 120 inhibitory synapses. The vertical dotted line indicates the normal parameter value used in all other simulations ($Ca^{2+}$, $K_V^+$, and $HCN_2$ conductances are only applicable for the active model).

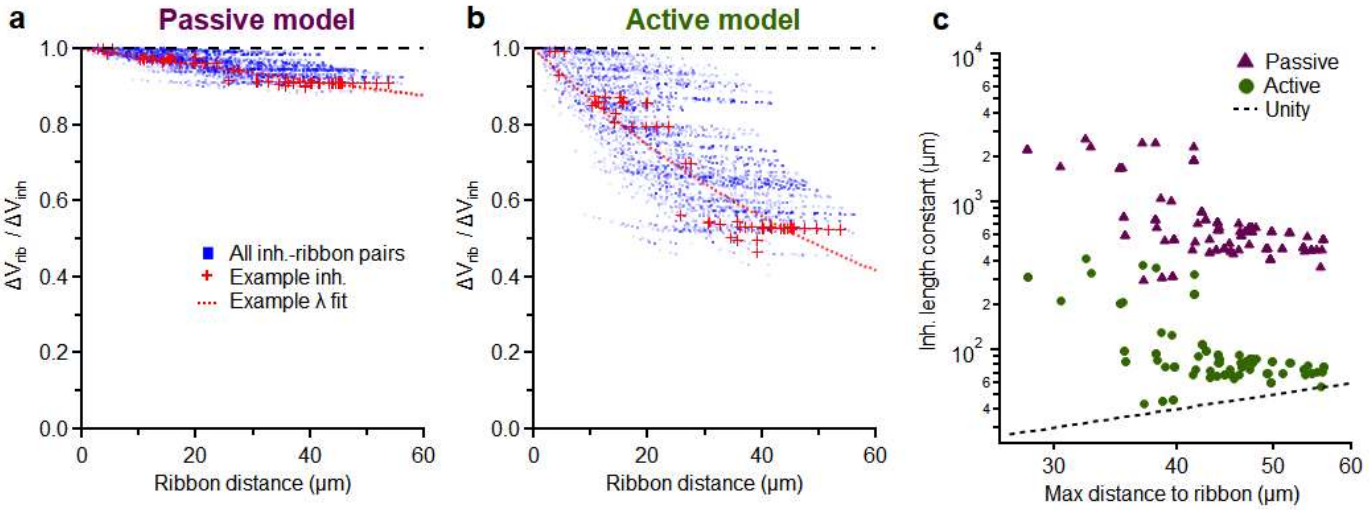


**Supplementary Fig. 10 | Fitting length constants to inhibitory synapses**
**a,b**, Cable model simulations in which a single inhibitory synapse is activated (same as **Fig. 6c**). The voltage change induced by inhibition is measured at the activated inhibitory synapse ($\Delta V_{inh}$) and at each ribbon output synapse ($\Delta V_{rib}$). The inhibitory voltage decay ($\Delta V_{rib}/\Delta V_{inh}$) is calculated for each ribbon output synapse and plotted against the path distance from the activated inhibitory synapse to that ribbon. Length constants are fit for each synapse according to the formula $\Delta V_{rib} / \Delta V_{inh} = e^{-x/\lambda}$, where x is the path distance and λ is the length constant. Length constants were calculated for both the passive (**a**) and active (**b**) versions of the type 6 BC model. **c**, The length constant of each inhibitory synapse plotted against the distance to the furthest ribbon from that same inhibitory synapse.
